# Deep Blueprint: A Literature Review and Guide to Automated Image Classification for Ecologists

**DOI:** 10.1101/2025.11.03.686223

**Authors:** Chloe A. Game, Nils Piechaud, Kerry L. Howell

## Abstract

1. Deep learning (DL) is a powerful tool to extract ecological information from large image datasets efficiently and consistently. However, applying these methods remains challenging, due in part to the complexity of DL workflows and the dynamic nature of available tools.
2. To address this, we created a practical guide and review, focused on one of the fundamental tasks in automated image analysis: image classification. Our approach integrates commonly used software and highlights key steps - from image acquisition to annotated, model-ready datasets, to training, evaluation and deployment. It is modular and supported by a flexible code base (Python & R) and Graphical User Interfaces (GUIs), enabling adaptation to different models and ecological objectives. The goal is to empower ecologists to confidently incorporate computer vision into their research.
3. We illustrate this approach, using an open-source ROV dataset from the Norwegian Sea, featuring deep-sea biotopes defined by multivariate clusters of depth, substrate type, and associated species. To balance accessibility for users alongside performance, we focused on CNN models from the Ultralytics ML Platform (YOLO V.8 and V.11), comparing the full suite of architectures that range in complexity and efficiency.
4. Cross-validation revealed high overall performances and that larger, more complex models are not always superior, with YOLO V.8m best (Accuracy = ∼0.98). Notably, high performances were achieved despite labels being based on both visual and external environmental predictors, suggesting visual features alone were sufficient for classification in this dataset. We highlight that the decision to deploy a model must be made in light of the study’s objectives, with domain-based reasoning and experience guiding every stage of implementation.
5. This work offers a practical blueprint for implementing DL in ecological research, promoting broader adoption and supports reproducibility and more efficient, standardized, and sustainable monitoring; in this case of deep-sea biotopes, which is essential for marine spatial planning.

## Introduction

As the need for knowledge-based conservation policies is increasingly acknowledged by scientists and biodiversity managers (Rogers *et al*., 2023), the development and improvement of sampling technologies is accelerating. A broad aim of these advancements is to substantially increase data gathering capacity for benthic ecologists (Brandt *et al*., 2016; Clark, Consalvey and Rowden, 2016; Levin *et al*., 2019). Many types of data are gathered, including oceanographic, acoustic, geological and chemical, to give an increasingly comprehensive representation of natural phenomena in or around the seafloor. For understanding the spatio-temporal dynamics of epibenthic megafauna and benthic communities, optical imagery has proved to be an effective tool (Howell, Davies and Narayanaswamy, 2010; Taylor *et al*., 2017) and now represents a large proportion of the data collected at the seafloor in the 21st century (Solan *et al*., 2003; Bicknell *et al*., 2016; Gomes-Pereira *et al*., 2016a). In the deep-ocean, this has largely been facilitated by the introduction of robotic platforms with imaging capabilities such as Remotely Operated Vehicles (ROV) (Ewing, Vine and Worzel, 1946; Ewing, 1967; Brandt *et al*., 2016; Clark, Consalvey and Rowden, 2016) and Autonomous Underwater Vehicles (AUV) (Morris *et al*., 2014; Wynn *et al*., 2014; Huvenne *et al*., 2018).

However, the increased rate at which ecologists can acquire images has not kept pace with their ability to analyse them, and this has led to what is termed an *image analysis bottleneck* (Schoening *et al*., 2017). This problem may not be unique to ecologists, but it is particularly amplified in deep-sea ecology given the insufficient number of scientists available to process image data compared to the spatial extent of the study area and the amount of data necessary to accurately represent this ecosystem. For ecologists, this processing largely involves manual annotation of imagery, in which image information is interpreted, or translated, to a semantic level (Gomes-Pereira *et al*., 2016a); an inherently slow task (Beijbom *et al*., 2012). Thus this limits the scale of image acquisition, or creates long delays before results are produced.

As a result, automated analysis of images with *Machine Learning* (ML) is increasingly being used to measure biological phenomena from large datasets (Katija *et al*., 2022; Belcher *et al*., 2023; Crosby *et al*., 2023). Convolutional Neural Networks (CNNs) (Lecun *et al*., 1998; Krizhevsky, Sutskever and Hinton, 2012; LeCun, Bengio and Hinton, 2015; Simonyan and Zisserman, 2015; Goodfellow, Bengio and Courville, 2016; He *et al*., 2016; Huang *et al*., 2017; Alzubaidi *et al*., 2021; Jocher, Chaurasia and Qiu, 2023), are widely popular ML algorithms in automated image analysis, including ecological applications (Weinstein, 2018; Rubbens et al., 2023). Their complex, or deep, model structure, with multiple layers that process image information to extract patterns and learn relationships, motivated their categorisation into a new subfield of ML called *Deep Learning* (DL) (Hinton and Salakhutdinov, 2006; LeCun, Bengio and Hinton, 2015).

In the last 5 years, there have been a number of marine applications using DL for classification of imagery, including classification of benthic taxa and habitats (Piechaud *et al*., 2019; Mahmood, Ana Giraldo Ospina, *et al*., 2020; Durden *et al*., 2021; Yamada *et al*., 2022; Jackett *et al*., 2023; Game, Thompson and Finlayson, 2024), plankton (in flow-cam imagery) (Kerr *et al*., 2020) and fish (Marini *et al*., 2018; Ditria *et al*., 2021; Villon *et al*., 2021). For further applications, see (Katija *et al*., 2021; Abad- Uribarren *et al*., 2022; Crosby *et al*., 2023; Rubbens *et al*., 2023). DL constitutes an important part of *Computer Vision* (CV), by which computers can be used to mimic the human visual system, processing and interpreting visual information. Employing such methods, in image analysis pipelines, is important to optimise existing approaches by promoting efficiency, consistency and quality of ecological data extraction (Cuvelier, Zurowietz and Nattkemper, 2024). However, whilst these benefits receive more and more recognition from both researchers and the media, knowing how to harness the underlying methodologies to answer biological questions is far from straightforward (Tuia *et al*., 2022). In addition, there is a clear need for better performances and efficiency (Pichler and Hartig, 2023; Misiuk and Brown, 2024) for CV-based solutions to compete with manual ones. Hence, many scientists and developers are constantly seeking to improve and innovate in this sector. While this is positive, one of the consequences is that the landscape is very dynamic and, like in the *red queen* ecological theory (Van Valen, Leigh, 1973), users must constantly keep up with innovation, with little time to digest a given method before it is superseded by a new one.

These challenges have led to the development of smoother interfaces between DL models and user annotations (Borremans *et al*., 2024; Clark *et al*., 2024) and, indeed, some image annotation softwares now possess this functionality such as BIIGLE (Langenkämper *et al*., 2017a), VIAME (Dawkins *et al*., 2017a), CoralNet (Chen *et al*., 2021), Tator (Tator, 2025), CVAT (CVAT, 2025) or Biodoc (Biodock, 2025). Some of the steps commonly found in ML workflows - mainly creating annotations, training and deploying models - can be performed in these softwares with their intuitive *User Interface* (UI) but not with the flexibility, consistency and speed offered by a programming interface. Thus, it is more practical and covers more use cases to script one’s implementation of ML workflows, particularly at scale (Belcher *et al*., 2023; Crosby *et al*., 2023).

There are many benefits to ecologists implementing machine learning workflows themselves, as it allows them to better understand the goals and nuances of their study, and to more effectively integrate their subject-matter expertise into both the workflow design and the interpretation of results (Game, Thompson and Finlayson, 2024). Interfaces that bypass the need for computer-science and programming skills exist and let ecologists focus on the ecological objectives of their study (Clark *et al*., 2024). Efforts to avoid engaging with programming might stem from the misconception that such skills are out of reach and are limited to ML applications. In this regard, a *demystification*, as termed by Belcher *et al*. (2023), could help give more ecologists the confidence that these tasks are within their reach, especially for those already familiar with programming-based methods for statistics, modelling, data management and visualisation, or Geographical Information Systems (GIS). In addition, automated image analysis is becoming increasingly simple to run and evaluate with minimal programming, particularly using default parameters for classic (widely-utilised) and even some state-of-the-art (SOTA) models (Jocher, Chaurasia and Qiu, 2023; Skorch, 2025). Yet it remains on the fringe of commonly used methods to gather ecological data in the deep sea. Thus there is a need to better illustrate implementation of these methods by a non-specialist, not to the degree of scientists working in ML, but as a practical implementation (Game, Thompson and Finlayson, 2024).

In this guide, we provide a flexible workflow that can be adapted to users’ needs and innovations in ML. This is presented with a case study of a typical benthic dataset of deep-sea biotopes, collected by an ROV in the Norwegian Sea (Meyer *et al*., 2023). Biotopes can be defined as a habitat (e.g. depth range and dominant substratum) and its associated species (Olenin and Ducrotoy, 2006; Costello, 2009). We approach the workflow holistically, connecting software and resources commonly used by deep-sea ecologists and explain steps involved from collecting image data to deriving an automatically interpreted output ready for ecological analyses. This work will encourage the regular usage of ML to gather ecological data in the deep sea, moving towards standardised, efficient and sustainable monitoring. Whilst primarily tailored to marine ecologists, this guide will be useful for beginners in other fields.

### Guided workflow and literature review

Here we present a guided example of an entire image classification pipeline, with relevant peer-reviewed literature and technical documentation such as code repositories and tutorials. This serves to simplify the process for novice users by highlighting six main tasks: (1) Objective, (2) Data preparation, (3) Annotation, (4) ML, (5) Testing & Deployment and (6) Deliverable. For further clarity, an overview of these steps is presented in Figure 1. Although each task is discussed in the ecological context of our case study, the broad methodological steps can be applied to alternative use cases. The methodology presented is, for the most part, conducted programmatically in Python (Van Rossum and Drake, 2009). This simplifies the pipeline into a single interface, offers more flexibility and, crucially, is transferable to alternative workflows and requires less manual input. A tutorial of this can be found at (Game and Piechaud, 2025), which can be used both locally (e.g. in a Jupyter Notebook, as in this study) and on the cloud to leverage public image data and computing resources (Google Colab). However, the results of the pipeline presented can also be achieved with a combination of standalone software and manual processing.

**Figure 1.**
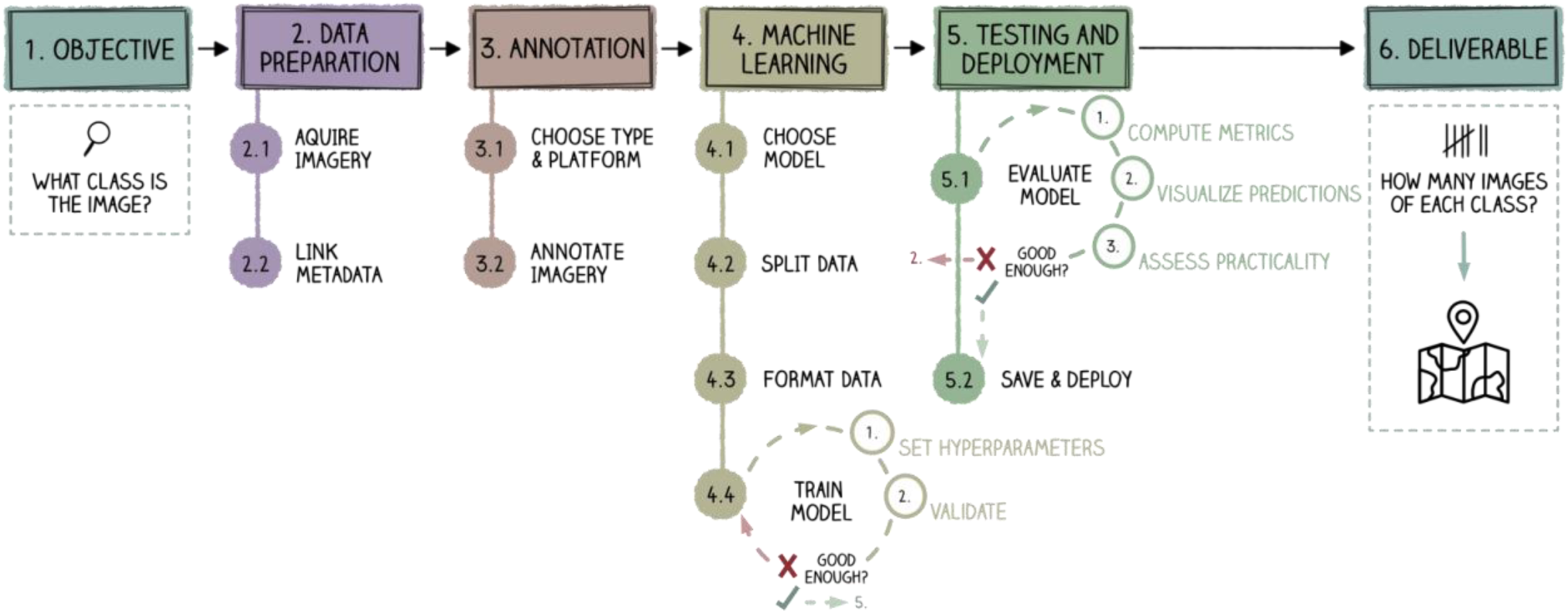
Simplified and idealized diagram of an image classification scenario. Each box represents a key task and corresponds to a section of this paper to aid comprehension. While presented largely linearly for clarity, real-world ML workflows are often iterative and non-linear and the need to revisit specific sections may vary depending on the scenario.

Most ML libraries are written in Python, therefore the primary code shared along with this study is in Python. Whilst many ecologists favour R (R Core Team, 2013), it is currently impossible to avoid Python completely in this workflow. To support accessibility, we provide examples of R code, including a shiny app for selected tasks. Note that Large Language Models (LLMs), such as ChatGPT (OpenAI, 2025), can support Python to R code translation, though they should be used cautiously, particularly with little understanding of the outputs.

## 1. Objective: What is your ecological task?

Before attempting any automated image analysis with ML, it is necessary to consider what ecological questions you wish to answer (Weinstein, 2018; Blair *et al*., 2024; Trotter, Griffiths and Whittle, 2025). Whilst this may seem obvious, these questions will affect your entire workflow. Only with a hypothesis in mind can you identify an appropriate model, prepare a suitable dataset and design a model training and evaluation procedure that has the potential to meet your objectives. Although we aim to provide ‘short-cuts’, building a ML pipeline is a significant investment in time and resources, requiring specific data types, hardware and some familiarity with programming and modelling concepts. Time and effort can be wasted if misinformed expectations are not met. Careful consideration and planning at the start of a project can avert such loss and help to decide if and how to engage with this method.

Within the scope of automated image analysis, broadly speaking there are three common ML problems; *Classification*, *Object Detection* and *Segmentation*. Classification, will identify, or ‘classify’, what each image represents i.e. assigning a category to each (Jackett *et al*., 2023; Trotter, Griffiths and Whittle, 2025). Methodologically speaking, the question is the same, whether the image is of a species or, as in this case, a deep-sea biotope. Note that in ML, the category we wish to assign to an object (or image) is referred to as a *class*. Should we wish to extract more detail, such as the location and abundance of a species within an image, the problem is referred to as Object Detection (Redmon *et al*., 2016). In one step, this both localises and classifies, ideally all, individuals of your target species (or class), with a box around the boundary of each individual. Furthermore, the spatial extent or size of a species (e.g. percentage cover) can be provided through Segmentation (Bearman *et al*., 2016). This ascertains which pixels of the image represent each class, producing a polygon, or box, around each pixel group found. Each of these ML problems are *image annotation* tasks and can be applied to multiple classes at a time. At their root, they are all classification problems. To use and understand them, is to first use and understand classification algorithms.

**Example 1 : Study Objective**

In our example, we wish to estimate the abundance, diversity and richness of multiple deep-sea biotopes in ROV imagery from Schulz Bank on the Arctic Mid-Ocean Ridge (Meyer *et al*., 2023), producing maps of their distribution, to contextualise their ecology in the surrounding environment and provide data to support marine spatial planning. This requires imaging a large amount of ground to statistically significantly represent the diversity of habitat. However we cannot realistically analyse this by hand, it must be performed automatically.

Classification of benthic images into biotopes remains one of the main tasks for which benthic scientists are required to provide expertise (e.g. Buhl-Mortensen *et al*., 2020; Ross *et al*., 2023) and a field of application that is poised to greatly accelerate with the implementation of multiple new technologies, including automated image analysis (Misiuk and Brown, 2024).

## 2. Data preparation

### 2.1. Acquire imagery

It is common to use existing manually annotated imagery to train ML models, however such annotations may have been informed by more than just the visible content, for instance expert knowledge or contextual understanding that the model cannot access (e.g. environmental, temporal or behavioural cues). This can lead to confusion and poor performance when training. Thus, when possible, it is better to acquire data for the purpose of ML, though, like any modelling, the relevance and quality of this data will greatly affect performance.

In benthic image analysis, it is well documented that the means with which an image is captured can have a very important impact on their interpretation during manual annotation (Beisiegel *et al*., 2017; de Mendonça and Metaxas, 2021) as it shows varying perspectives of the seabed. File type and resolution of imagery may also affect the detection of small fauna/objects, faunal density estimates and the taxonomic resolution of the classifications (Schoening *et al*., 2020; de Mendonça and Metaxas, 2021). Thus the suitability of the imaging platform must be carefully aligned with the objectives of the ML task as it not only determines what can be captured, but also shapes the overall quality of the dataset.

There are a number of characteristics that contribute to data quality; one of the most important being consistency. ML models struggle with variability in the appearance of images and their content (e.g. taxa), with respect to characteristics such as the colour, orientation and size. Aside from some natural biological variation, these are tightly linked to the imaging platform used (e.g. AUV, ROV) and whether there is variation in its altitude, angle (incidence & tilt), field-of-view and lighting configuration, augmenting the direction and intensity for example. Since ML models must find and describe relationships within the data to make a prediction, increased fluctuation or noise makes it even more challenging to extract a clear signal from complex image data. Thus it is desirable to standardise imagery in this regard. That being said, if you hope to use a model on new datasets with their own, likely disparate, set of characteristics, too little diversity risks overfitting the model to the training data and may lead to poor generalisation. Variation in imaging platform can also constitute a *domain shift* for automated annotation, meaning the data distribution differs from what the model was trained on, potentially degrading model performance (Beery, Van Horn and Perona, 2018; Schneider *et al*., 2020; Katija *et al*., 2022; Belcher *et al*., 2023).

ML model performance is also sensitive to image degradation effects linked to light attenuation and scattering, such as reduced contrast and colour intensity as well as blurring and colour-casts (e.g. blue/green tint). Furthermore, the increased requirement for artificial lighting with depth often produces non-uniform illumination (e.g vignetting, halos and shadows) in imagery. Imagery may be obscured by equipment or water turbidity, induced by suspended sediment and marine snow for example. In the event that images are extracted video stills (frame grabs), be aware that images may also contain artefacts if the video codec has used interlacing to optimise the size of a video. All these factors compromise visual quality, distorting and obscuring discriminative taxonomic features (Durden *et al*., 2016; Schoening *et al*., 2020), which can in turn lower model performances.

Removing poor quality images is an obvious way to avoid ambiguous annotations biasing the models during training. Jackett *et al*., (2023) observed that carefully filtering noisy images from training data enhanced model accuracy by as much as 12%, while also reducing the computing load. However this could overfit the model towards ideal conditions which are less representative of ‘real-world’ data.

Another important contributor to dataset quality is the size i.e. number of images. DL models require large datasets to learn effectively and avoid overfitting, particularly in complex classification tasks. Model performance is well known to be influenced by the quantity of images representing each class, as this helps the model capture the full range of conditions it may encounter during deployment (Piechuad et al., 2019; Dawson, Dubrule and John, 2023; Venkataraju *et al*., 2023; Blair *et al*., 2024). This is a challenge in the marine domain, where datasets rarely cover all relevant conditions. Consequently, users may be forced to aggregate data from multiple sources, necessitating a commonly accepted annotation scheme, see Section 3. The lack of standardisation complicates such efforts. It is difficult to give precise guidance on a suitable dataset size, as it is both task- and data-dependent, and there is a diminishing return to adding more and more images. However, reasonable performances have been reported within magnitudes of 100s-1000s training images for each class (Piechaud *et al*., 2019; Jackett *et al*., 2023; Game, Thompson and Finlayson, 2024; Vega *et al*., 2024a).

To limit data quality issues, care must thus be taken at the acquisition stage. You should, where possible, collect a dataset that is consistent, yet appropriately diverse for your problem, large enough to support learning, ideally with a balanced class representation, no duplication and good visibility of your target classes. Images should also be relevant, representative and captured at an appropriate resolution to capture discriminative taxonomic features. However, often scientists must make-do with poor or varying data quality due to uncontrollable events, for example during fieldwork. Whilst some post-processing strategies can help such as image normalisation (colour/brightness), enhancement (Lisani *et al*., 2022) and augmentation (size, orientation), see Section 4.3, they cannot fully compensate for poor input data, and may render ML infeasible, necessitating a return to manual annotation. Decisions in the image acquisition phase should thus be grounded in the study’s objectives and a detailed understanding of the data, rather than relying on blanket rules alone.

### 2.2. Link Metadata

Although information about the image dataset, such as where, when and how it was collected, can be extremely useful for automated (and manual) image classification (Yamada, Prügel-Bennett and Thornton, 2021), here it is not fundamental to the learning process, which focuses on image content only (pixel information). Nonetheless, image metadata like latitude, longitude, depth, height above seabed, camera zoom, angle and speed can all be useful in pre-processing data, such as extracting images from specific regions and depths or investigating and improving the consistency and quality of the dataset. It can also be useful in the interpretation and interrogation phase, investigating whether there are mistakes that correspond to any metadata for example. Hypothetically speaking, this could help to distinguish why certain images are correctly identified as a soft coral habitat in one set of images but not in another set of images. Perhaps in the latter set the altitude of the camera was increased, augmenting the appearance of the habitat relative to the training data and reducing the visibility of taxonomic details that the ML model has associated with that class.

Aside from being generally good practice to collect image metadata in image analysis pipelines, if ML is to be better integrated, it may be necessary in order to deploy a suite of models that are optimised to specific metadata attributes such as region, depth or platform. Moreover, it can future-proof datasets, facilitating the development and adoption of emerging ML approaches, such as multi-modal transformers (Radford *et al*., 2021; Yu *et al*., 2022), especially since it is difficult to predict what information may prove valuable over time.

## 3. Annotation

Classification requires imagery to be annotated, or labelled, with a class name (semantic label) (Gomes-Pereira *et al*., 2016b) to train a ML model to associate the image content with a single class. Thus the training data could simply be a manually-generated data file (e.g. .csv, .txt or .xlsx) containing a table of dimensions *m* (rows) x *n* (columns), where *m* = number of images and *n =* 2, with one column for image filename and one for the assigned class. Alternatively, image annotation can be undertaken in specialised software, either public or private, as is common.

### 3.1 Choose platform

Annotation softwares typically enable efficient collaborative manual annotation and facilitate standardisation of data processing and annotation protocols, including usage of common labels (e.g. taxonomic or geological (Althaus *et al*., 2015; Howell *et al*., 2019a; European Environment Agency, 2022). A number of studies have attempted to make a census of public annotation softwares, showing how wide the choice currently is. ZenML (2025) for example, lists 28 options for images alone, which do not include the most popular tools for marine benthic imagery. To our knowledge, the list of those relevant for benthic imagery (Gomes-Pereira *et al*., 2016b; Langenkämper *et al*., 2017a; Bowden *et al*., 2020) lack recent updates.

Hence, there is no single best tool. The priority with any of these choices is that the software is actively maintained and that annotation data can be easily imported and exported. However, there are a number of other benefits, including improving the transparency and accessibility of image analysis, a quality considered most important for reproducibility in marine imaging (Borremans *et al*., 2024). Their usage will also facilitate the transition to object detection and/or segmentation methods at a later stage, which require complex annotation formats that cannot be achieved through manual data entry (Belcher *et al*., 2023).

The utility of annotation softwares continues to grow with the integration of automated annotation methods (Dawkins *et al*., 2017b; Zurowietz *et al*., 2018; Chen *et al*., 2021) and some support visualisation and tracking of ML outputs by enabling the upload of predicted labels (e.g. (Howell *et al*., 2023)). With the implementation of APIs, the integration of the different steps of the proposed workflow (Figure 1) can be scripted to an even greater extent, saving time and reducing potential manual errors. This could for example enable verification of model predictions, help to interpret model behaviour and identify and correct anomalies such as mislabelled images.

### 3.2 Annotate imagery

Assigning a class to an image is, in principle, a straightforward task. However, the choice of labels and annotation protocol should be guided by the objectives of the study and carefully defined before annotation begins. Numerous classification schemes have been developed to serve different purposes such as biodiversity management, risk evaluation or exploitation suitability (Strong *et al*., 2019). Others focus on standardized taxonomic and/or geological descriptions such as SMarTaR-ID (Howell *et al*., 2019b), CATAMI (Althaus *et al*., 2015) and EUNIS (European Environment Agency, 2022). These schemes provide predefined labels that may be used for *supervised classification*, where the model is trained to recognise classes assigned by the user; provided these class labels can be consistently and reliably assigned to images without additional information. In contrast, *unsupervised classification* automatically annotates images based on the inherent structure of the dataset (e.g. clustering). Whilst this may better reflect the variability within a specific dataset, it limits comparability to other studies (Fraschetti *et al*., 2024) and may not be meaningful for the project end-goal .

Biases stemming from poor data quality can emerge as early as the annotation stage. Minimising human error is therefore a crucial consideration when annotating imagery. Prioritising quality of annotations, rather than maximising volume, is more effective in producing high-performing models, even though it may require more manual effort (Catalán *et al*., 2023; Jackett *et al*., 2023; Prior *et al*., 2023). To support this, robust QC procedures and clear annotation protocols are essential, particularly when multiple annotators contribute to the same dataset (Culverhouse *et al*., 2003, 2014; Curtis *et al*., 2024; Cuvelier, Zurowietz and Nattkemper, 2024; Fraschetti *et al*., 2024; Pavoni *et al*., 2024).

Despite the recognised importance of consistency, there is no universally-accepted protocol for generating globally compatible datasets, although multiple studies have outlined best practices for annotating images for ML (Golding, N., *et al*., 2021; Catalán *et al*., 2023; Prior *et al*., 2023; Rädsch *et al*., 2023; Zhang *et al*., 2023). Standardisation across individual annotators and teams, and over time, is vital, not only for broader marine ecology research (Howell *et al*., 2019a), but also to ensure ML models can learn from clear and reliable patterns. Inconsistent labelling between annotators can obscure these patterns (Trotter, Griffiths and Whittle, 2025). Variation and discrepancies in file formats or archiving routines also present additional challenges for data integration and reuse (Howell *et al*., 2019a; Katija *et al*., 2021; Schoening *et al*., 2022; Dawson, Dubrule and John, 2023). Annotations should thus follow a consistent format, classification scheme and taxonomic precision to support both reproducibility and clarity (Howell *et al*., 2019a; Moore *et al*., 2019; Borremans *et al*., 2024).

**Example 3.2 : Dataset**

In our case, we use a small open-source dataset of the Schulz Bank seamount on the Arctic Mid-Ocean Ridge (Meyer *et al*., 2022). This contained 600 RGB images (1500x930 *.png*), collected by an ROV, that were classified into 5 novel Biotopes: AB, C, K, S & X, see Figure 2. Importantly, such biotope descriptions are critical for marine spatial planning, since they are not captured by existing official classification schemes such as EUNIS (European Environment Agency, 2022). Although these biotope labels are abstract, compared to something more widely intuitive such as a *Hard-bottom Sponge Aggregation* or *Deep-Sea Seapen Community* for example, they provide essential descriptions of megabenthic communities.

Annotating the images with these biotopes, differed to the standard annotation procedure for image classification, in which a domain expert inspects each image and assigns a class label. Using BIIGLE 2.0 (Langenkämper *et al*., 2017b), key species were localized within each image, using point annotations, and the dominant substrate was recorded. Then a hierarchical cluster analysis of the present species, dominant substrate and depth, in PRIMER-E 7.0 (Clarke and Gorley, 2015), was used to derive the biotopes. Each biotope in the dataset is thus a multi-modal classification, representing a multivariate cluster of depth, dominant substrate and associated species. Further methodological details are provided in Meyer et al. (2023).

## 4. Machine Learning

### 4.1. Choose model

At its simplest, classification with ML involves learning a relationship between some input variable(s) and output classes (encoded as integer labels). With image classification, each pixel value can be considered an input variable, however this is typically not sufficient information to enable accurate classification. Thus a traditional approach is to extract image characteristics linked to colour, textural and geometric patterns (Haralick, Shanmugam and Dinstein, 1973; Lowe, 2004; Matas *et al*., 2004; Dalal and Triggs, 2005; Mikolajczyk and Schmid, 2005; Ahonen, Hadid and Pietikainen, 2006; Bay, Tuytelaars and Van Gool, 2006) and classify with algorithms such as Support Vector Machines (Cortes and Vapnik, 1995), K-Nearest Neighbours (Guo *et al*., 2003) or Decision Trees (Quinlan, 1986; Ho, 1995). Such hand-crafted image features have been used in marine applications, aiding classification of corals (Stokes and Deane, 2009; Beijbom *et al*., 2012), fish (Kitasato *et al*., 2018; Tharwat *et al*., 2018), kelp forests (Mahmood, Ana G. Ospina, *et al*., 2020) and other marine megafauna (Lopez-Vazquez *et al*., 2020). However, these methods do not typically generalise well across datasets (Eerola *et al*., 2024). Alternatively, CNNs have become very popular having surpassed these performances (Lopez-Vazquez *et al*., 2020). These deep networks have the advantage in that, during training, they design custom image filters to extract features that improve classification accuracy. For details on CNN training, see Section 4.4.

**Figure 2.**
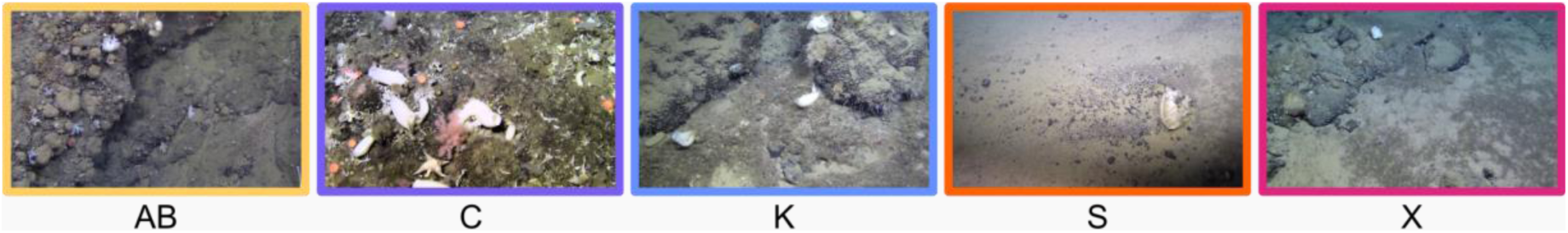
Examples of the Schulz Bank biotope classes used in this study.

Broadly speaking, CNNs have a widespread reputation for excelling in image classification tasks (Maurício, Domingues and Bernardino, 2023; Mienye *et al*., 2024). There are a large number of CNN architectures and models from which to choose and given the fast pace of CV research, new architectures are seemingly released every day. Commonly employed CNN architectures in image classification include ResNet (He *et al*., 2016), VGG (Simonyan and Zisserman, 2015), MobileNet V2 (Sandler *et al*., 2019), DenseNet (Huang *et al*., 2017) and Efficient Net (Tan and Le, 2019). This is thanks to their notable performance on benchmark classification datasets such as ImageNet (Deng *et al*., 2009), that act as a standardized measure of their accuracy. Many of the CNNs that have been developed are modifications of these ‘baseline’ architectures and aim to incrementally improve performance, speed or generalizability.

Another popular group of CNN architectures, which are widely adopted in CV (particularly object detection) are variants of the You Only Look Once (YOLO) architecture (Redmon *et al*., 2016). These models have become increasingly applied to marine ecological research (Fayaz, Parah and Qureshi, 2022).

More recently, Vision Transformers (ViTs) have gained momentum as an alternative to CNNs, yet they often incur higher computational costs and require larger training datasets to avoid overfitting (Dosovitskiy *et al*., 2021). Though approaches such as DeiT (Touvron *et al*., 2021) are helping to mitigate these limitations. Given these constraints and the extensive documentation and long-established success of CNNs, particularly in resource-limited settings, they offer a more practical, entry-level choice in this work.

A consequence of the high diversity and abundance of CNN architectures is that there is no universally-accepted best solution and the final choice for a given project will have to compromise between factors such as performance (on related tasks), computing costs, compatibility with available hardware, ease of use and deployment or familiarity among users. SOTA models for example, may achieve top accuracies on benchmark datasets but can be challenging for non-specialists to implement. Particularly when newly released models may not yet be supported, if at all, within the most commonly adopted and user-friendly frameworks such as Pytorch (Paszke *et al*., 2019; PyTorch, 2025), Ultralytics (Ultralytics, 2025a) or TensorFlow (Abadi *et al*., 2016; TensorFlow, 2025). These frameworks typically incorporate models that are popular and thus familiar amongst peers - a useful feature when disseminating work and encouraging uptake. Employing models within these frameworks will be simpler due to standardized coding approaches that generalize across architectures. In addition, they often provide more comprehensive documentation and are more likely to receive long-term support, should licensing permit. This long-term support, from sustained developer involvement or an active community of users, is especially important for large or long-term projects, where ongoing maintenance, error resolution and updates are needed to keep pace with evolving standards and hardware.

Other considerations in model choice include usability, interoperability and scalability. Is the model and its associated tools straightforward to understand and use, reducing the learning curve for new users? Will the results be compatible with existing workflows or evaluation standards, ensuring that outputs can be readily interpreted, compared, or integrated into broader analytical pipelines? Can the model handle increasing data and complexity without significant performance bottlenecks?

Data and computational requirements are particularly critical factors in selecting a model. Deeper and more complex CNN architectures (e.g., ConvNeXt (Liu *et al*., 2022; Woo *et al*., 2023), ResNet (He *et al*., 2016), ResNeXt (Xie *et al*., 2017)) have a high capacity to capture intricate patterns in imagery. However, this comes with more parameters, increasing memory demands, and may require larger datasets to perform well. They also may require more *Floating-Point Operations* (*FLOPs*), for example additions and multiplications, which increase the computational cost. Together, these factors may lead to slower training times and require more powerful servers or cloud infrastructure to operate efficiently in future deployment. Whereas more efficient architectures (e.g. EfficientNets (Tan and Le, 2019, 2021), MobileNets (Howard *et al*., 2019)) are designed with fewer parameters and FLOPs. Whilst this may sacrifice some accuracy in difficult tasks, they require less data, enable faster training and can run on more constrained hardware. Although ultimately, this trade-off is nuanced and context dependent.

Importantly, data and computational demands can be alleviated with *Transfer Learning,* in which a model is pre-trained on a larger dataset (e.g. ImageNet) to recognise general image features such as edges, shapes and textures. This model is then trained further (fine-tuned) on a new, often smaller, dataset to recognise task-relevant features. Although the classification task differs from the pre- training, this often leads to faster training and improved performances and is thus employed in many ML workflows. Transfer learning can make a training dataset of 100s or even a few 1000s of images viable (D’souza, Huang and Yeh, 2020; Villon *et al*., 2021; Venkataraju *et al*., 2023), whilst magnitudes more may be needed to train from scratch (Deng *et al*., 2009; Krizhevsky, 2009; Kuznetsova *et al*., 2020). Scratch-based DL is therefore often too big an endeavour for a small scale project with limited data, time and computational resources.

**Example 4.1 : Model Choice**

In this work, accessibility was the primary concern in our model choice. We thus initially selected the YOLO (Redmon *et al*., 2016) family of architectures, since their integration into the Ultralytics ML platform (Ultralytics, 2025a) has made it user-friendly and accessible to non-expert users.

In simple terms, Ultralytics is a high-level wrapper around PyTorch, offering a collection of Python functions that streamline common tasks in the ML workflow, such as training, inference (prediction), evaluation (including visualisations), and deployment. It allows users to create a workflow with basic knowledge of ML concepts and there are a number of documented code examples and videos on how to get started, both from Ultralytics and the large community of YOLO users. For simple tasks such as making predictions with an existing model, this is simpler than alternative or more advanced frameworks such as Keras (Keras, 2025) or Pytorch. However, with more complex tasks, the learning curve remains steep.

The relative homogeneity of the Ultralytics framework has made it possible to transition between classification, object detection, segmentation and tracking tasks with ease, bringing many possibilities to the ML beginner. For classification specifically, Ultralytics only enables users to train and deploy the native YOLO models (versions 8 & 11). To ensure a more comprehensive evaluation, we compare the whole suite of YOLO V.8 architectures, which range in size and complexity (number of trainable parameters), from the smallest V.8n to the largest V.8x (2.7-57.4 million parameters). We also evaluate the newer YOLO V.11 architectures, which are computationally lighter (<28.4 million parameters) and improve on the YOLO V.8 benchmarks (Ultralytics, 2025a). In a transfer learning step, we adapt each of these models, which are pre-trained on ImageNet, to the Schulz Bank dataset.

Both YOLO V.8 & V.11 are designed for efficiency and ease of use, achieving accuracies between 70 and 80% on the ImageNet-1k dataset (Ultralytics, 2025a). In comparison, top-performing CNN classifiers, EfficientNetV2-L (Tan and Le, 2021) and ConvNeXt V2-H (Woo *et al*., 2023) achieved 85.7% & 86.3% accuracy, respectively, but at a much higher cost, with 120 million & 660 million parameters. Thus, while these YOLO models do not achieve the highest accuracy, their relatively small-size, straightforward implementation and integration within a well-documented library that also supports other CV tasks, makes them a practical and accessible choice. However, they are by no means the only possible way to achieve the goals of this example study and users should build their workflow with modularity in mind to facilitate uptake of better tools as they become available.

### 4.2. Split data

In order to train and evaluate a DL model effectively, a separate training, validation and test dataset is required, though these are typically derived by partitioning a single dataset (Xu and Goodacre, 2018). Intuitively, the training dataset is used to train, or teach, the model to perform a task through machine learning (LeCun, Bengio and Hinton, 2015). The validation dataset is used to validate, or assess the performance of, the model during training. This will influence choices made by the researcher when developing a model and guide internal modification of model parameters during training, to improve the ability of a model to perform a task. Therefore a separate test dataset, not involved in training or validation, is necessary to assess the final model performance, once training has finished. This process of splitting data and testing the model on unseen data is referred to, in ML terms, as *hold-out validation*. It is an extremely important technique to minimise overfitting, wherein a model performs very well on training data but generalises poorly to independent data - a largely undesirable model behaviour in ML. Importantly, for hold-out validation to truly lessen overfitting, validation and test data must be independent of the training data. Validation and test images must therefore not occur, nor correlate to, the training set images, for example by featuring duplicate or overlapping images which contain the same individual taxon. This is a form of *data leakage* which undermines the reliability of the model and can lead to inflated performance.

Data sets are commonly split into 60-80% training, 10-20% validation and 10-20% testing data, for example (80, 10, 10) or (70, 15, 15). However, it is well understood in ML that a training set only needs to be large enough to represent the variability of the target data, rather than choosing an arbitrary proportion. The same is true for the validation and test sets; additional images will have little effect on performance, provided the data is representative and offers a reliable estimate of generalisation (Xu and Goodacre, 2018; Durden *et al*., 2021).

Another common method to partition data is using *K-fold Cross Validation* (Kohavi, 1995). Here the data is first split into training & testing (hold-out validation), then the training dataset further split into *k* folds (groups), with *k*-1 folds used for training and the remainder used as a validation set, for example where *k*=5, 4 splits (80%) are used for training and 1 split (20%) used for validation. This is a more comprehensive technique that better accounts for stochasticity in the data, giving a general view of performance that is not specific to any one data split. It is more robust to overfitting and gives a measure of the variability one can expect from performance of models originating from different splits.

Many datasets, including that used in this study, suffer from an imbalanced number of classes. Some classes will be naturally more prevalent than others or are perhaps captured more easily by the optical imaging platform used. In these cases, it may be desirable to stratify the data partitions, enforcing the same class ratios across partitions. This will be important to train a model for a class distribution that occurs within the unseen testing data. Although stratified splits can be achieved manually (Piechaud *et al*., 2019), it is more efficient and repeatable to do this programmatically. This is true regardless of the approach chosen to split the data, however it is important to use a seed for traceability. This is a number that will ensure that any random split of data can be repeated exactly, ensuring reproducibility of results. Note that data partitions are often physically separated into digital folders (e.g. train, validation & test) in ML workflows, including YOLO V.8 & V.11. However, some, such as those employing Pytorch models and functions, allow images to be tracked with indexing and handle splits internally.

**Example 4.2 : Data Partition Strategy**

In this work, all data was stored and processed locally. We split the dataset into train and test partitions of 80% and 20% respectively and employed a class-stratified *k*-fold cross validation of the training set, where *k*=5. All splits were seeded.

### 4.3. Format data

Before training a DL model, input data must be formatted according to the architecture and training data (in the case of transfer learning). In some workflows (e.g. YOLO) these steps are handled automatically, however in the interest of clarity we detail potential formatting steps below.

Typically, input variables to a CNN, here the image pixel values, should be small to support training stability and performance. It is therefore standard to normalise them between 0-1. If using a pre-trained model it may also be necessary to standardise your input data to that which the model was trained on. By ensuring that the input data has a consistent scale and distribution to the original training data, it will help preserve model performance and speed up processing and convergence. A typical approach is to center and scale the image pixel values, by subtracting the mean RGB value of the training set and dividing by the standard deviation (Brownlee, 2020b).

It is also often necessary to resize images to a size expected by the architecture, for example 224 x 224 pixels (Russakovsky *et al*., 2015; Simonyan and Zisserman, 2015; He *et al*., 2016; Howard *et al*., 2017). Images can first be resized along the longest dimension, maintaining the aspect ratio to prevent distortion. However, since images are typically not square, their spatial dimensions (height and width) should also be adjusted. This reshaping can be achieved by cropping (e.g. center cropping) or padding (e.g. with zero values). Note that the depth dimension, or colour channels (R, G & B), are preserved (e.g. 224 x 224 x 3). The appropriateness of these methods will depend on the task.

Given a choice of image size, as is the case with YOLO V.8 & V.11, as a rule of thumb, the highest resolution is usually best for preserving image details that distinguish the classes and give the model more pixels to work with when improving its accuracy. However, this comes with added computational costs, which may exceed hardware capacities, and negatively impact both training and inference speed. In practice, higher image resolutions tend to result in better performances but not systematically (Liu, Brailsford and Bull, 2025). Users must therefore consider the trade-off between accuracy and feasibility. Starting with downsized images and gradually increasing the resolution during model development is a practical way to appreciate the coupled performance gains and speed losses that come with increasing resolution.

Although not required, augmentation is another important formatting tool, typically applied prior to training, though test-time augmentation strategies also exist (Luz *et al*., 2025). Augmentation involves systematically modifying images, simulating additional variability, to better capture unseen characteristics within your dataset. Common strategies include geometric (e.g. rotation, flipping, scaling) or photometric adjustments (e.g. brightness, contrast or colour changes). This can lead to improved model performances and generalisation (Zoph *et al*., 2020; Song *et al*., 2022; Sun, Yue and Li, 2022; Belcher *et al*., 2023) without incurring additional computationally-driven costs to inference, unlike model architecture complexification (Zoph *et al*., 2020). However it is a complex topic and can have variable effects on different classes within the same model (Belcher *et al*., 2023). It is also domain-specific and may have unpredictable effects on the model and subsequent use of it. For example, the generic augmentation methods known to improve models on primarily land/air imagery can degrade performance of models trained on underwater imagery (Tan, Langenkämper and Nattkemper, 2022). We recommend careful consideration before implementing any augmentation and to review this in light of the data, task and deployment environment.

**Example 4.3 : Data Formatting**

All default YOLO formatting was accepted in this work except image size which was set to 256 pixels along the longest edge. The choice was made to optimize performance while remaining close to the scale of 224, a common image size in other classification architectures. It is important to note that 224 is incompatible with YOLO’s architecture, as dimensions must be multiples of 32 for spatial downsampling.

For simplicity, we also chose to accept the default augmentations that are applied during training. Although it can yield great benefit of time and resources to carry out the search for beneficial augmentations (Tan, Langenkämper and Nattkemper, 2022)

### 4.4. Train model

In DL, training a CNN is a trial and error process. Simply put, a sample (or *batch* in ML) of training images are passed to the model, which extracts features (pixel patterns) using learnable parameters called *weights* (e.g. image filters) and *biases*. Note that in scratch-based learning, these parameters are initialized randomly, in contrast to transfer learning, which starts with pre-trained parameters that are already partially optimized.

Extracted features are used by the model to make predictions about which class each image belongs to. This step, in which image data is passed forward through the CNN, is referred to as *Forward Propagation* (Goodfellow, Bengio and Courville, 2016). The resulting classification is a set of probability scores (returned by the *softmax* function), a proxy for confidence, that the image represents each class (e.g Class 1= 0.6, Class 2= 0.3, Class 3= 0.1). These are then compared to the ground-truth (e.g. Class 1 = 1, Class 2= 0, Class 3= 0) and the error, a type of *loss*, is measured.

This error is used to adjust the model parameters responsible for feature extraction and classification (weights & biases), aiming to improve the performance on the next batch of images. With the degree to which they are adjusted, typically proportional to the size of the error; a model making large errors will need to be modified more than one making smaller errors. These updates happen by first determining the contribution of each parameter to the error, sequentially in a backward direction through the CNN, in a step termed *Back Propagation*. The parameters are then updated using an optimization algorithm such as *Stochastic Gradient Descent.* For explanation on these algorithms, refer to (Starmer, 2022).

Once all training image batches have passed through the CNN, completing one *epoch*, the model enters a validation stage; classifying the withheld validation images and calculating the loss. By selecting the highest softmax score (*argmax*), accuracy can also be calculated, as the proportion of images assigned the correct classification i.e. the classification assigned during manual annotation. The training and validation process iteratively enhances model performance and continues until some condition is met, for example a maximum number of epochs.

#### 4.4.1 Set hyperparameters

Training a CNN requires a number of hyperparameters to configure the learning process, however these are not optimized during model training and so must be tuned manually by the end-user. Importantly, this can result in significant gains in performances. However, since typical optimization strategies involve the training of multiple models to find the best hyperparameter combinations, for example an ablation study or grid search, this is an extremely time-consuming and resource-hungry process. End-users will thus have to evaluate this trade-off in the context of their own ML problem.

Some crucial hyperparameters to the training process include the *learning rate* which, relative to the loss, scales the degree of parameter adjustment during training, influencing how sensitively the model responds to mistakes. Choosing this value is challenging and has led to the development of *learning rate optimizers* such as Adam (Kingma and Ba, 2015), that dynamically alter this value during training to improve efficiency and speed.

The *loss function* can also be adjusted according to the task. For multi-class classification, categorical cross-entropy is the standard choice, helping the model to be more confident in the correct predictions, or highest probabilities. Alternatives include sparse categorical cross-entropy and Kullback-Leibler divergence loss or for binary classification, binary cross-entropy (default), hinge loss or squared hinge loss (Brownlee, 2020a).

Considering the number of images passed to the model at once (*batch size*) is also important, with links to training speed, resource usage and performance, and requires balancing trade-offs. Small batches may slow training and cause instability, but can improve generalization by introducing noise; large batches train faster and produce more stable updates, but require more memory and may lead to overfitting (Keskar *et al*., 2017).

The number of epochs also plays an important role in efficiency and performance, as it controls how long the model learns from the data. Too few epochs can lead to underfitting, whilst too many can lead to overfitting. An arbitrary number can be chosen and training intercepted once performance stabilizes. However, a more robust and resource-efficient approach is to employ *early stopping*; Here model training stops when the performance has not improved over a set number of epochs e.g. the validation loss has not reduced.

Each of these hyperparameters will have a default and thus the training process can be reasonably straightforward, albeit not optimized to each classification task. We therefore recommend optimizing the batch size and number of epochs, as well as employing early stopping, for a simple, yet computationally light way to see improved performances. Note that the tuning options presented are not exhaustive; the range of possible adjustments is too vast to comprehensively cover in one paper and there are likely to be more as the field advances. For further hyperparameter and training configuration details, please refer to (Ultralytics, 2025a).

**Example 4.4.1 : Training Hardware and Hyperparameter Selection**

Training was possible using a NVIDIA RTX A100 6GB Laptop GPU, given the small size of the dataset. Note that larger datasets may require GPUs with higher compute powers to avoid significant delays to training speed and memory errors.

All default hyperparameters were accepted except batch size, number of epochs and early stopping for which we chose 32, 50 and 5, respectively. Upon completion of training, the epoch model with the best validation performance was chosen as our final trained model. For each YOLO V.8 & V.11 architecture, this resulted in 𝑘=5 models, which were automatically saved by YOLO.

#### 4.4.2 Validate

Model quality can be assessed from several angles, both during and post-training. Typical metrics include measures of loss, accuracy and speed as well as visualizations such as *confusion matrices*, all of which we discuss in more detail below.

Although not commonly reported, monitoring the behaviour of the loss function throughout training is essential to effectively assess the model’s ability to learn or generalize. Since this is an error metric that should be minimized, the training and validation loss should decrease over epochs, producing an ‘l-shaped’ distribution, as evidenced in Figure 3.

**Figure 3.**
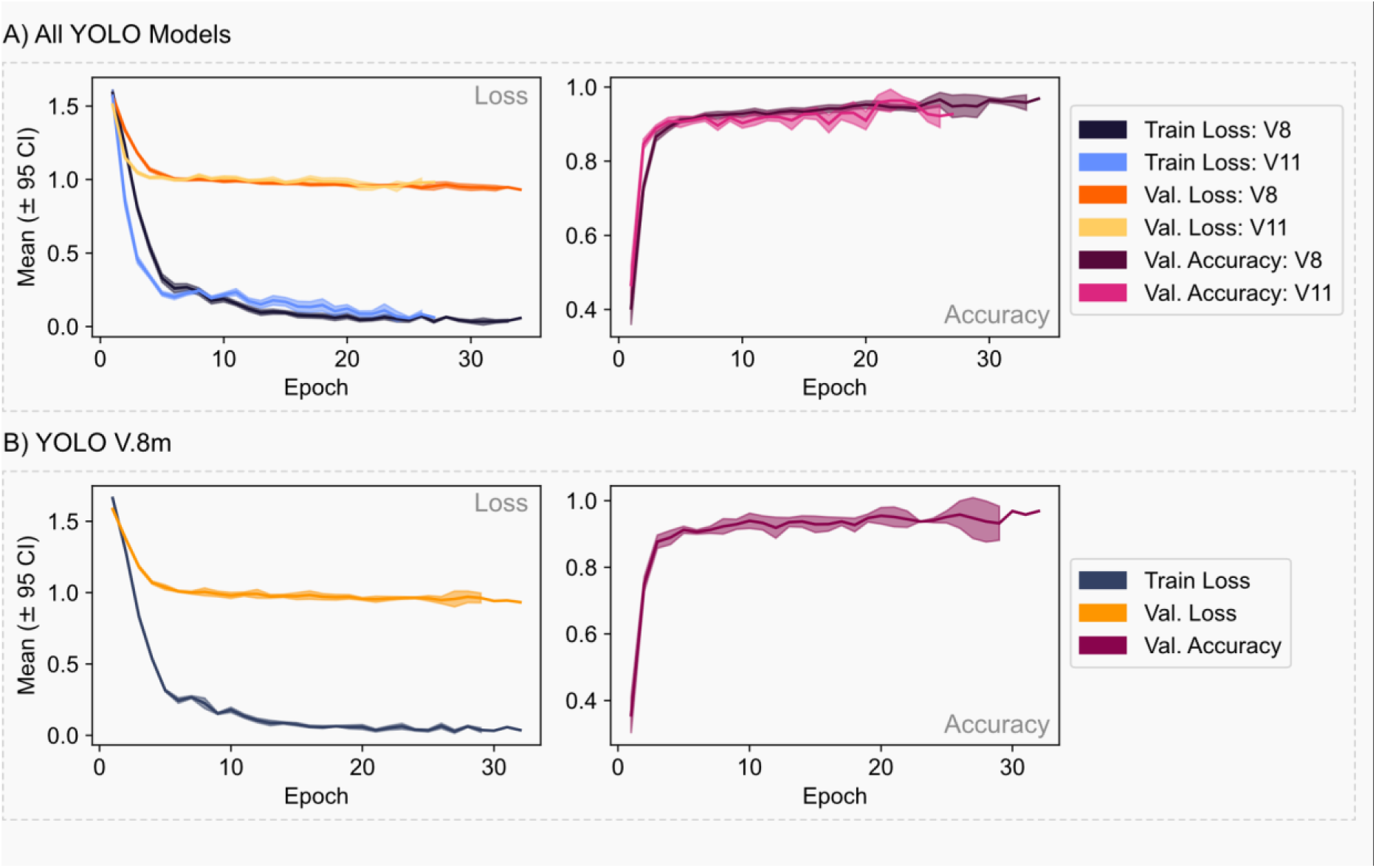
Mean epoch training and validation performance across k-folds for (A) all YOLO models (V.8 vs V.11) and (B) YOLO V.8m

Aside from loss, a typical, and perhaps the simplest, way to measure model performance is to determine the accuracy (often referred to as *Top-1 Accuracy*)*. Top-5 Accuracy* may also be used, where a prediction is considered correct if the true class is among the five highest-scoring predicted classes. This can be a useful indicator of model performance and uncertainty in classification problems with many classes, such as ImageNet-1k (1000 classes). However, this has no merit in few-class classification tasks, such as the 6-class task in this study.

**Example 4.4.2 (a) : Epoch-wise Training and Validation Assessment**

YOLO V.8 models were found to have lower, and more consistent, training and validation loss than V.11 on average (Figure 3a). However, both exhibited steady declines over training epochs. Rapid early learning was indicated by a sharp loss decrease and spike in validation accuracy in the first five epochs, after which point the loss broadly plateaued. However the gentle increase in validation accuracy until early-stopping, suggested the models continued to learn useful class-discriminative features despite no improvements to model confidence.

For all models, the training loss was significantly, and largely consistently, lower than the validation loss, which could suggest a model that is overfitting to the training data. However, given the complexity of marine datasets such as this, in which the classes are often highly imbalanced, have few images and exhibit high intra-class variation, it is expected that performance on the validation data will suffer. Despite this, Figure 3a. shows that mean validation accuracy (Top-1) peaked around 0.96 for both YOLO V.8 & V.11 models, at epochs 34 and 22 respectively, indicating effective generalization.

Epoch training and validation performance for YOLO V.8m is also presented in Figure. x3b for completeness. Whilst the epoch analysis did not indicate a significant advantage of this architecture, V.8m ultimately achieved the best cross-validation performance (see remainder of Section 4.4.2), supporting its selection as the preferred architecture for further analysis.

Upon completion of training, it is also beneficial to analyse the predictive performance of the final models in more detail. Accuracy is again an important metric, however this does not reveal any information on the relative performance of each class so additional metrics are required, namely *recall*, *precision* and *f1-score*. These metrics are commonly used in test set evaluation (Section 5.1.1) and, like accuracy, all return a score between 0 & 1, with 1 indicating *perfect* performance.

Rec**all**, or sensitivity, measures the proportion of ground truth images of a given class that the model correctly identifies i.e. of **all** the images of class 1, how many were classified as class 1 by the model. **Pre**cision, on the other hand, evaluates the accuracy of the model’s predictions for a given class i.e. for all images **pre**dicted as class 1, how many were actually class 1 in the ground truth.

Depending on your task, the importance of these metrics to evaluate model performance may be weighted differently. For example, if identifying all images of class 1 is more important, even at the cost of over-predicting its occurrence and lowering precision, then the recall should be favoured. Alternatively, if ensuring predicted presences of class 1 are correct, at the expense of missed occurrences and lowering recall, then precision should be favoured. In cases where the two metrics are of equivalent value, the harmonic mean of the two, referred to as the f1 score, can also be insightful as a unique performance measure.

For each class 𝑐, performance metrics are calculated using Equations 1-3.

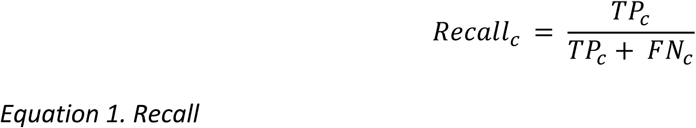

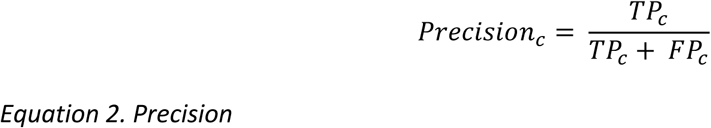

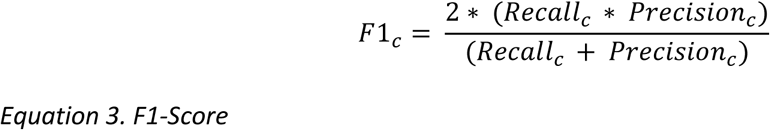

where 𝑇𝑃 (true positives) are images of class 𝑐 correctly classified as class 𝑐, 𝐹𝑃 (false positives) are images of other classes incorrectly classified as class 𝑐 and 𝐹𝑁 (false negatives) are images of class 𝑐 incorrectly classified as another class.

Note that these metrics can be aggregated by their mean performance across classes, often referred to as the macro-average. Since this weights class importance equally, it is useful with imbalanced datasets where rare or singleton classes are as important as the most abundant classes. The macro-average for each metric 𝑚 is calculated by Equation 4., where 𝐶 is the total number of classes.

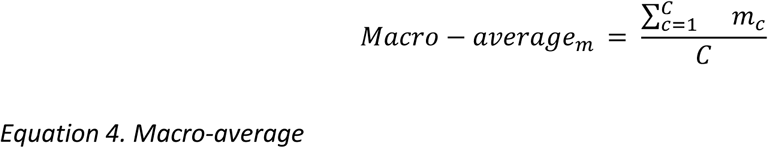

If the majority classes are more important, weighted averages will be more appropriate. Cohen’s Kappa & Matthews Correlation Coefficient may also be useful to assess the overall agreement and account for chance in imbalanced datasets.

**Example 4.4.2 (b) : Final Validation Performance**

Five final YOLO models were produced for each architecture, using the best epoch model in each *k*-fold. These models demonstrated strong overall performance, achieving average scores between 0.95 & 1 on the training set and between 0.93 & 0.97 on the validation set (Table 1). Class-performance metrics also indicated that the models were both accurate and well-balanced between precision and recall.

Comparing V.8 & V.11 models (Figure 4), performance decreased from training to validation on average, by 0.02-0.04, as expected from the epoch analysis (Example 4<u>.4.2a</u>). In all metrics, V.8 models slightly outperformed V.11, by 0.02 on the training set and 0.01 on the validation set.

Within the V.8 family, the larger architectures (8m, 8l & 8x) performed best, with highly consistent and near-perfect training performance (Table 1). Validation performance also remained high (>0.95), though greater variation was observed across class metrics; a characteristic seen across all architectures.

Among these V.8 models, YOLO V.8m narrowly emerged as the top performer, with mean *k*-fold metrics around 1 for training and between 0.96 and 0.97 for validation.

**Figure 4:**
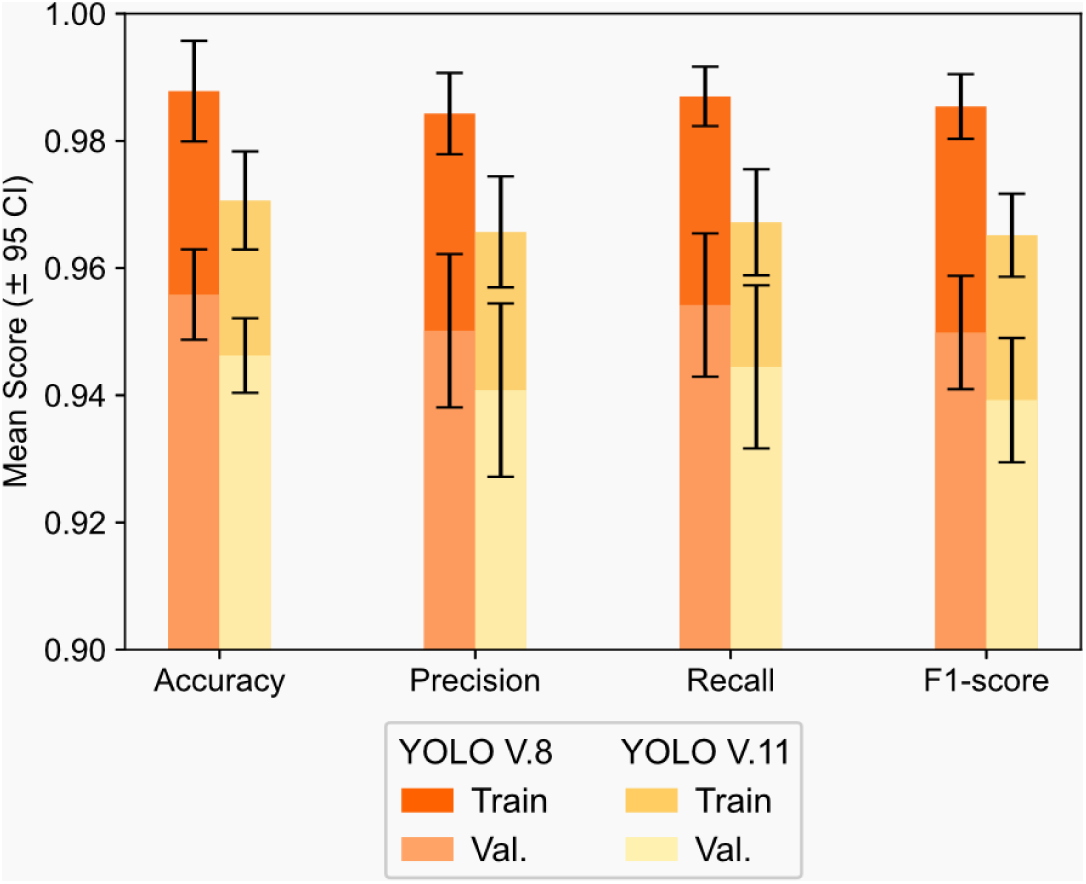
Mean training and validation performance for final YOLO V.8 & V.11 k-fold models

The values that form the basis for calculating the performance metrics (accuracy, recall, precision and f1-score) are also captured within *confusion matrices*, a standard diagnostic tool in ML (Figure 5).

**Figure 5.**
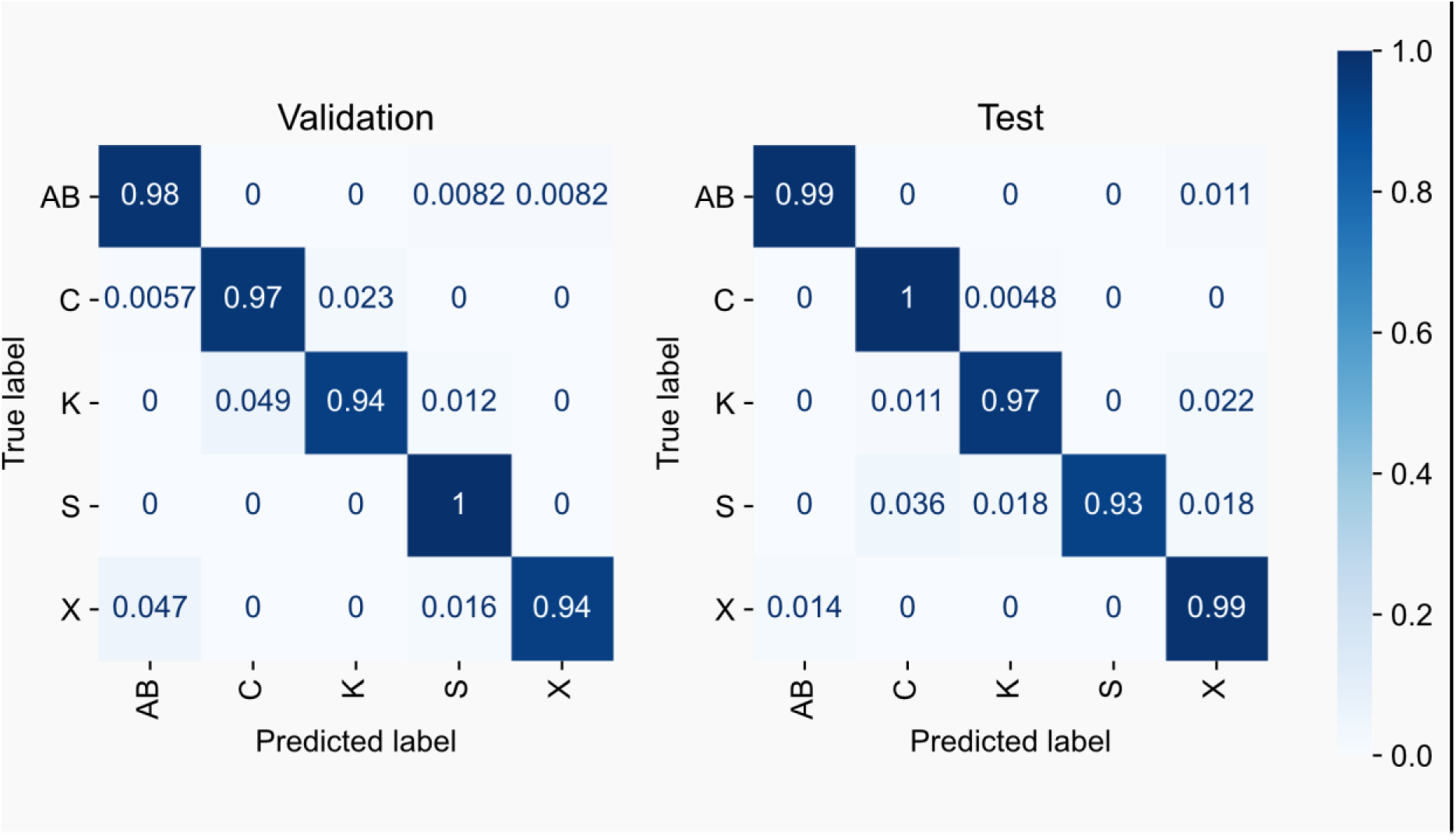
Summed and row-normalized k-fold matrices for Validation and Test Data.

**Table 1.** Training and validation performance: Mean k-fold scores (± standard deviation) on the training and validation sets for the final YOLO models.

| Model | Accuracy |  | Precision |  | Recall |  | F1-score |  |
| --- | --- | --- | --- | --- | --- | --- | --- | --- |
|  | Train | Val | Train | Val | Train | Val | Train | Val |
| **11n** | 0.97 (±<br>0.027) | 0.95 (±<br>0.023) | 0.97 (±<br>0.055) | 0.94 (±<br>0.079) | 0.97 (±<br>0.037) | 0.95 (±<br>0.073) | 0.97 (±<br>0.04) | 0.94 (±<br>0.056) |
| **8n** | 0.97 (±<br>0.023) | 0.94 (±<br>0.014) | 0.96 (±<br>0.041) | 0.93 (±<br>0.086) | 0.97 (±<br>0.034) | 0.94 (±<br>0.082) | 0.96 (±<br>0.033) | 0.93 (±<br>0.063) |
| **11s** | 0.97 (±<br>0.026) | 0.94 (±<br>0.016) | 0.96 (±<br>0.061) | 0.93 (±<br>0.088) | 0.96 (±<br>0.044) | 0.93 (±<br>0.09) | 0.96 (±<br>0.041) | 0.93 (±<br>0.071) |
| **8s** | 0.98 (±<br>0.032) | 0.96 (±<br>0.017) | 0.97 (±<br>0.061) | 0.95 (±<br>0.075) | 0.98 (±<br>0.039) | 0.95 (±<br>0.069) | 0.98 (±<br>0.046) | 0.95 (±<br>0.06) |
| **11m** | 0.98 (±<br>0.01) | 0.96 (±<br>0.005) | 0.98 (±<br>0.024) | 0.95 (±<br>0.055) | 0.98 (±<br>0.025) | 0.95 (±<br>0.069) | 0.98 (±<br>0.018) | 0.95 (±<br>0.044) |
| **11l** | 0.98 (±<br>0.015) | 0.95 (±<br>0.014) | 0.98 (±<br>0.037) | 0.95 (±<br>0.076) | 0.97 (±<br>0.067) | 0.94 (±<br>0.074) | 0.97 (±<br>0.041) | 0.94 (±<br>0.059) |
| **8m** | **1.0 (±**<br>**0.003)** | **0.97 (±**<br>**0.017)** | **1.0 (±**<br>**0.009)** | **0.96 (±**<br>**0.048)** | **1.0 (±**<br>**0.006)** | **0.97 (±**<br>**0.056)** | **1.0 (±**<br>**0.005)** | **0.96 (±**<br>**0.041)** |
| **11x** | 0.96 (±<br>0.011) | 0.94 (±<br>0.009) | 0.95 (±<br>0.058) | 0.94 (±<br>0.09) | 0.95 (±<br>0.052) | 0.94 (±<br>0.061) | 0.95 (±<br>0.035) | 0.94 (±<br>0.047) |
| **8l** | 0.99 (±<br>0.007) | 0.96 (±<br>0.021) | 0.99 (±<br>0.016) | **0.96 (±**<br>**0.059)** | 0.99 (±<br>0.014) | 0.96 (±<br>0.06) | 0.99 (±<br>0.01) | **0.96 (±**<br>**0.042)** |
| **8x** | **1.0 (±**<br>**0.002)** | 0.96 (±<br>0.012) | **1.0 (±**<br>**0.003)** | 0.95 (±<br>0.07) | **1.0 (±**<br>**0.003)** | 0.96 (±<br>0.05) | **1.0 (±**<br>**0.003)** | **0.96 (±**<br>**0.04)** |

1. Models are ordered by increasing parameter count. Bold font dictates best results.

These comparison tables of the validation ground truth and predictions, serve as a visual aid to identify performance patterns. They identify classes that are correctly identified and those mistaken for each other e.g. distinguishing between true/false positives and negatives. They can also highlight class imbalance effects, for example if a model performs poorly on rare classes. Confusion matrices are often normalized over the true labels (e.g. row-wise normalization), calculating the proportion of correct and incorrect predictions for each true class. A model with perfect performance would thus display a proportion of 1 (or 100% if a percentage) for each class down the center diagonal. Generating confusion matrices is simple with python libraries such as scikit-learn (scikit-learn, 2025), or in the case of YOLO (Ultralytics, 2025a), produced automatically following training.

**Example 4.4.2 (c) : Confusion Matrices**

In our study, since the validation confusion matrices varied extremely little over *k*-folds, we present the summed and row-normalized *k*-fold matrix for the top performing model, YOLO V.8m, in Figure 5. While some variation exists amongst classes, the confusion matrix indicates strong performance (>0.94) across both abundant and rare classes; an important quality for imbalance datasets like this.

Aside from predictive performance, monitoring speed *and memory requirements* is also an important evaluation of a model’s resource demands and deployability in real-world scenarios. In Ultralytics, these are automatically produced in the standard console output during training. Understanding the limitations, with respect to time or computational resources, helps to ensure that the model not only performs well but can also operate efficiently within the resource constraints of the target application.

**Example 4.4.2 (d) : Speed Analysis**

As demonstrated in Figure 6, training time increased somewhat with the number of trainable parameters, but remained generally fast, ranging from ∼2.5 to 4 minutes on average. Significant differences in training speed were indicated only between the smallest models (V.11n, V.8n, V.11s) and largest (V.8l, V.8x). However, at this dataset size, the benefits of faster training for smaller models was marginal. While YOLO V.11 models span a narrower set of parameter quantities than YOLO V.8, most YOLO V.8 models still fall within this range (<30M) and require a similar training time. Notably, this includes V.8m, the best performer in terms of accuracy and class performance metrics, showing that higher performance does not necessarily entail significantly longer training times.

**Figure 6.**
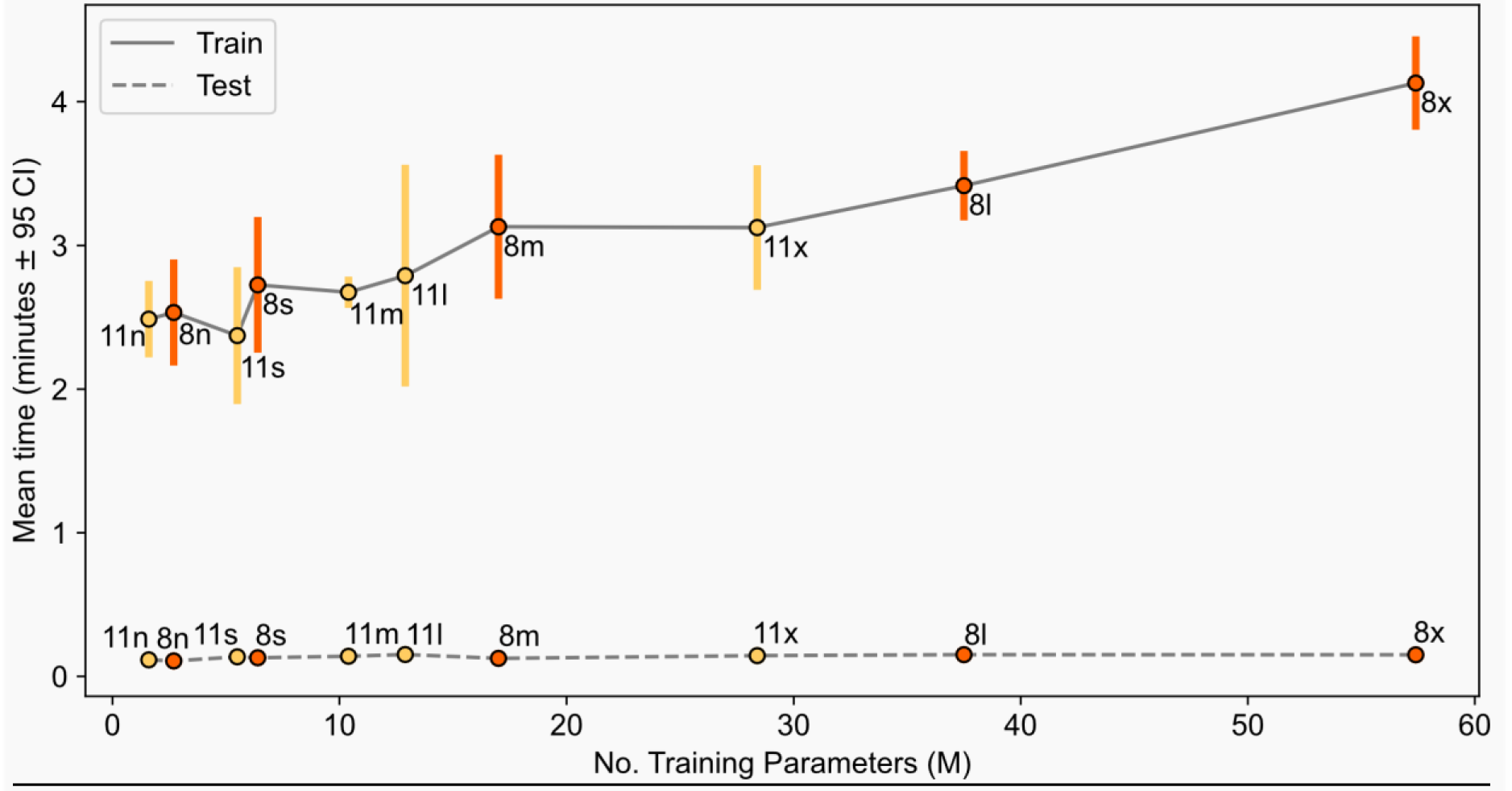
Mean training and inference time for all images in the training and test sets, respectively.

Following validation, if the model is not good enough you may have to return to earlier stages in the ML pipeline (Figure 1). ML tasks are rarely solved linearly. The challenge, however, is identifying the cause of the poor performances. Those driven by model structure or behaviour may require further tuning of the model hyperparameters (Section 4.4.1) or adjustments to model choice and data formatting (Sections <u>4.1</u> & <u>4.3</u>). Alternatively, you may have to address the data themselves, returning to the Data preparation and Annotation stages (Sections <u>2</u> & <u>3</u>).

**Example 4.4.2 (e) : Optimal Model Selection**

In our case, YOLO V.8m was chosen as the final model since it offered the optimal trade-off between training time, performance and stability. Given the high training and validation performance, the 5 final V.8m models proved to be ready for Testing and Deployment (Section 5), to evaluate performance on unseen data, designed to represent a real-world scenario.

Additionally, we continue some evaluation of alternative YOLO architectures to provide comparative insights (Figure 6).

## 5. Testing & Deployment

Although the final validation assessment (Section 4.4.2) provides a useful indication of model performance, it may yield overly optimistic results. This is because the validation data directly influences model development, through parameter tuning and early-stopping for example, introducing a risk of data leakage. While techniques like cross-validation help reduce dependency on any single validation subset and improve the robustness of model selection, they do not eliminate this risk. Thus we must evaluate the model on an independent hold-out test set. Although the test set is often still drawn from the same overall dataset, it offers a more reliable estimate of the model’s ability to generalise to truly unseen data.

Methodologically, this testing stage mirrors the final validation step, using the same inference and largely the same evaluation procedures. While the core evaluation metrics remain consistent, the testing phase may also include additional visual analyses and practicality assessments. Despite their similarities, we refer to this stage as *evaluation* to avoid confusion with the specific role *validation* plays earlier in the ML pipeline. The results of this evaluation will inform whether the model is ready for real-world deployment or requires further refinement. In the former case, the model can then be prepared accordingly, to ensure it is usable, reliable and maintainable in practice.

### 5.1. Evaluate model

#### 5.1.1 Compute metrics

Generalisation performance can be quantified with the standard evaluation metrics: accuracy, precision, recall and f1-score. Applying the same metrics as those used during validation provides a consistent basis for comparison, ensuring any score differences reflect true performance changes. It also will indicate whether the model is overfitting to the validation data (large discrepancy), or underfitting if both validation and test performance is poor. In the latter case, however, the model does typically not progress to the testing and deployment stage.

**Example 5.1.1 : Test Performance Metrics**

Consistent with the validation performance, the models generalized well to the unseen test data, achieving accuracy and macro-averaged class metrics between 0.91 & 0.98 (Table 2). The models also indicated a good balance between prediction quality and quantity, with similar macro-averaged precision and recall. These results are particularly positive given the small training set and that the class labels (biotopes) are based on both visual and external environmental predictors. This suggests visual features alone were sufficient for classification in this dataset, although external data could improve robustness in ambiguous cases and novel datasets (e.g. multi-modal learning). Despite some variation in training time, all YOLO models were highly efficient and consistent during inference, averaging only 8.4 seconds per fold *(SD=* 1.8; *n=*120) (Figure 6).

**Table 2.**
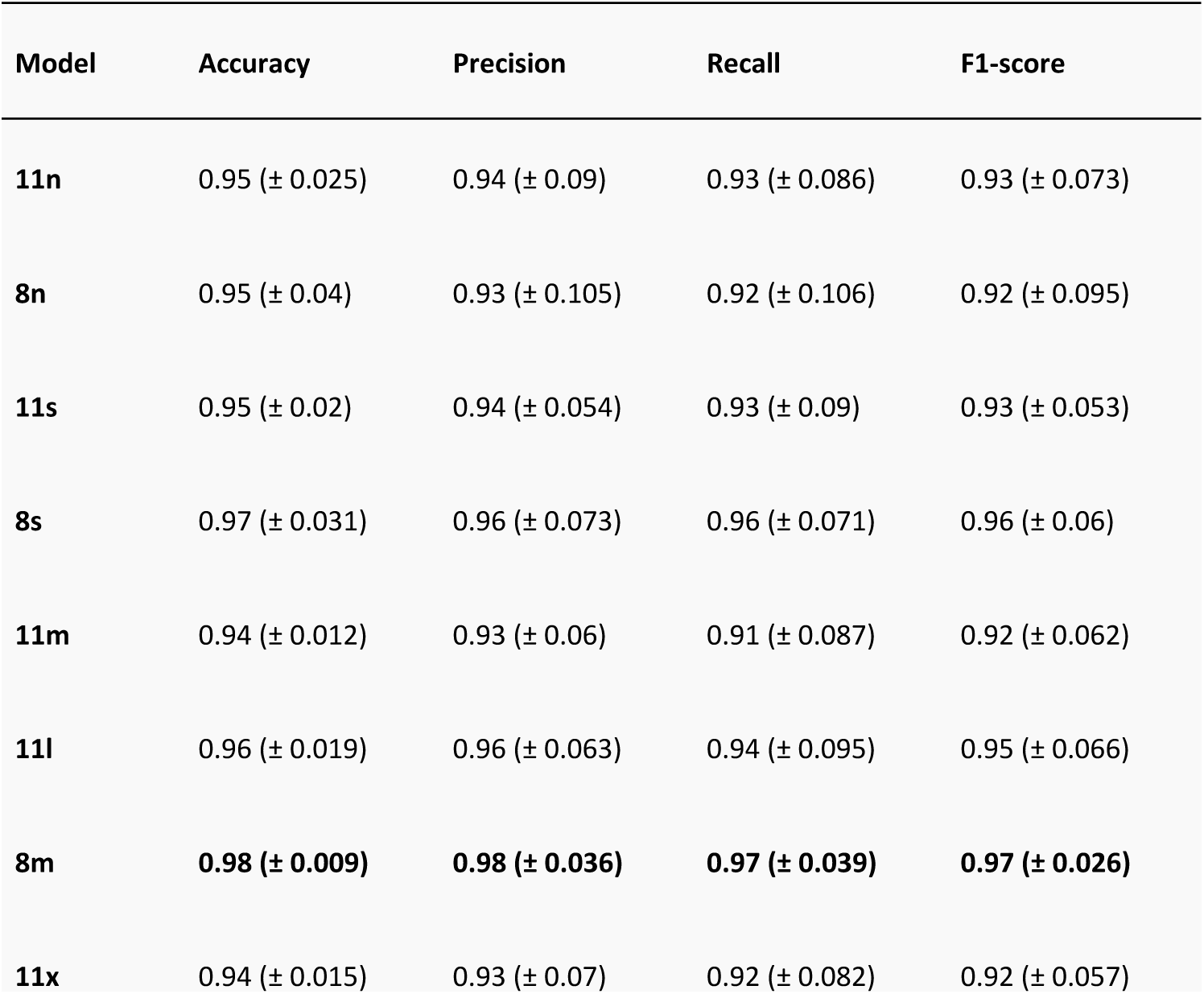
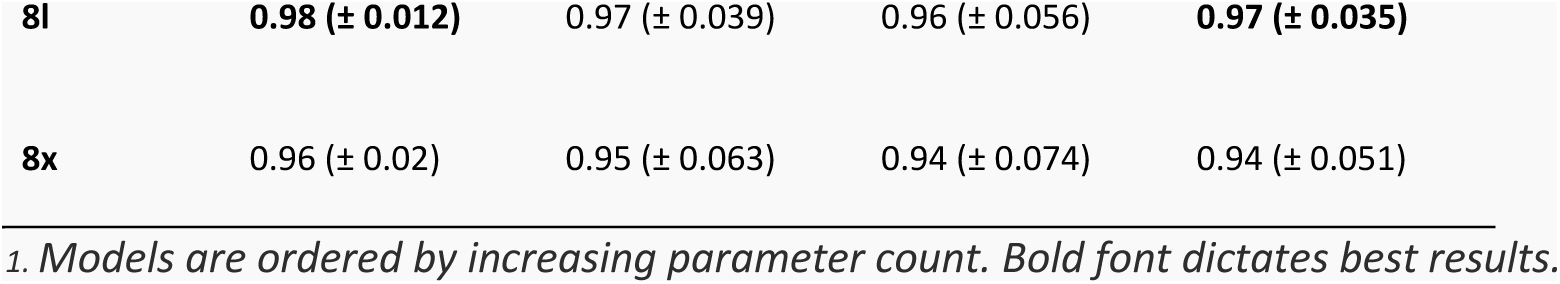
Test performance: Mean k-fold scores (± standard deviation) for all YOLO models.

| Model | Accuracy | Precision | Recall | F1-score |
| --- | --- | --- | --- | --- |
| 11n | 0.95 (± 0.025) | 0.94 (± 0.09) | 0.93 (± 0.086) | 0.93 (± 0.073) |
| 8n | 0.95 (± 0.04) | 0.93 (± 0.105) | 0.92 (± 0.106) | 0.92 (± 0.095) |
| 11s | 0.95 (± 0.02) | 0.94 (± 0.054) | 0.93 (± 0.09) | 0.93 (± 0.053) |
| 8s | 0.97 (± 0.031) | 0.96 (± 0.073) | 0.96 (± 0.071) | 0.96 (± 0.06) |
| 11m | 0.94 (± 0.012) | 0.93 (± 0.06) | 0.91 (± 0.087) | 0.92 (± 0.062) |
| 11l | 0.96 (± 0.019) | 0.96 (± 0.063) | 0.94 (± 0.095) | 0.95 (± 0.066) |
| 8m | **0.98 (± 0.009)** | **0.98 (± 0.036)** | **0.97 (± 0.039)** | **0.97 (± 0.026)** |
| 11x | 0.94 (± 0.015) | 0.93 (± 0.07) | 0.92 (± 0.082) | 0.92 (± 0.057) |
| **8l** | **0.98 ($\pm 0.012$)** | 0.97 ($\pm 0.039$) | 0.96 ($\pm 0.056$) | **0.97 ($\pm 0.035$)** |
| **8x** | 0.96 ($\pm 0.02$) | 0.95 ($\pm 0.063$) | 0.94 ($\pm 0.074$) | 0.94 ($\pm 0.051$) |

1. Models are ordered by increasing parameter count. Bold font dictates best results.

Comparing architectures, YOLO V.8 variants, except V.8n, consistently outperformed V.11 (Table 2). Though, on average, this improvement was marginal at 0.02 (2.4 images) for each metric (Figure 7). These findings demonstrate that newer models do not inherently guarantee better performance and further highlight the value of evaluating multiple architectures, should resources permit.

**Figure 7.**
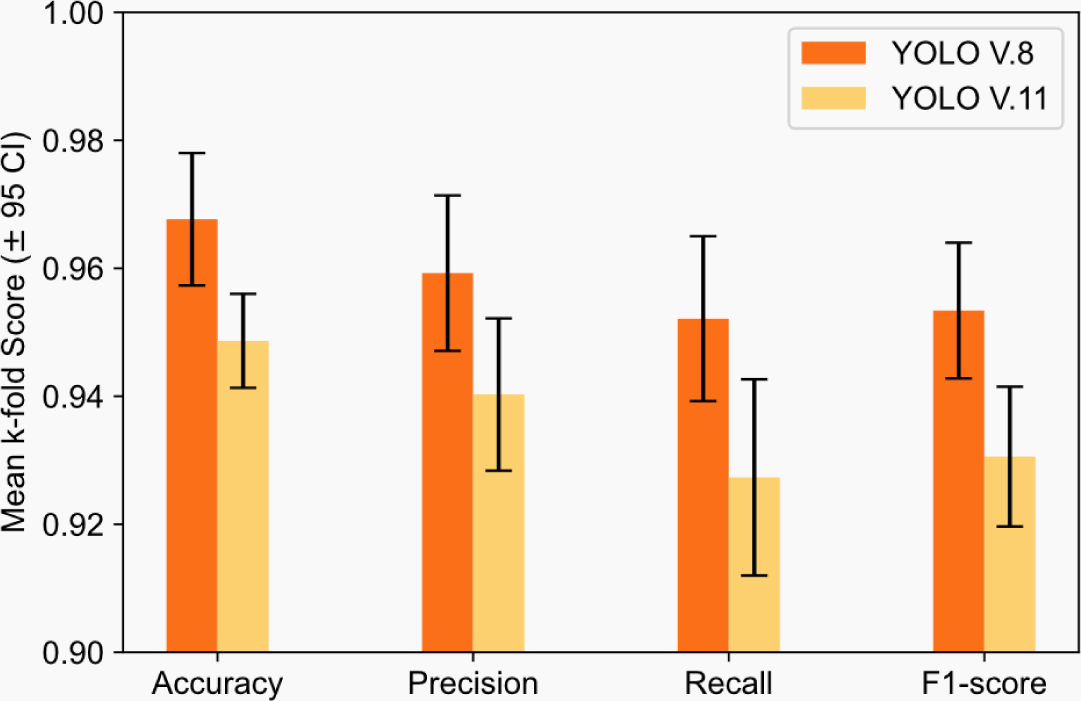
Test performance: Mean k-fold scores (± 95 confidence intervals) for YOLO V.8 & V.11 models.

Although all models performed well overall, the V.8m evaluation revealed a slight, but consistent advantage, reinforcing its selection as the optimal model during cross-validation stage (Section 4.4.2). Among the V.8 models, V.8m was consistently best (>0.97) (Table 2). While its performance gain over other *V.8 models* was small (≤0.05, or 6 images), these differences may become more pronounced with a larger dataset. Furthermore, V.8m exhibited the most stable performance across folds, with a standard deviation <0.04 across metrics.

Much like the validation stage, the strong classification performance of V.8m is well displayed in a confusion matrix, summed and normalized over *k*-folds (Figure 5). This showed that the model rarely misclassified images across classes. These few misclassifications, shown in more detail in Figure 8, tend to decrease with increasing class prevalence, an effect particularly reflected in the F1-scores.

Intuitively, the model was most confident predicting the most common training classes C & AB, with average softmax scores around 0.99 (Figure 8). This was coupled with the highest f1-scores (also ∼0.99), suggesting clear class separability, and more stable prediction metrics across folds. In comparison, the model was slightly less confident when predicting the rarer classes, X & K, with average confidence scores between 0.94-0.95 respectively. For class X, this uncertainty was reflected by the lower, yet still good, precision rate of 0.93. Despite their lower prevalence in the dataset, performance for both X & K was satisfactory, with F1 scores between 0.96 & 0.97. Interestingly, whilst the model was very confident in identifying the rarest class, S, and achieved perfect precision on average (1.00), its mean recall was the lowest amongst the biotope classes at 0.93 (Figure 5). Across the remaining classes, precision and recall were more balanced and consistent.

**Figure 8.**
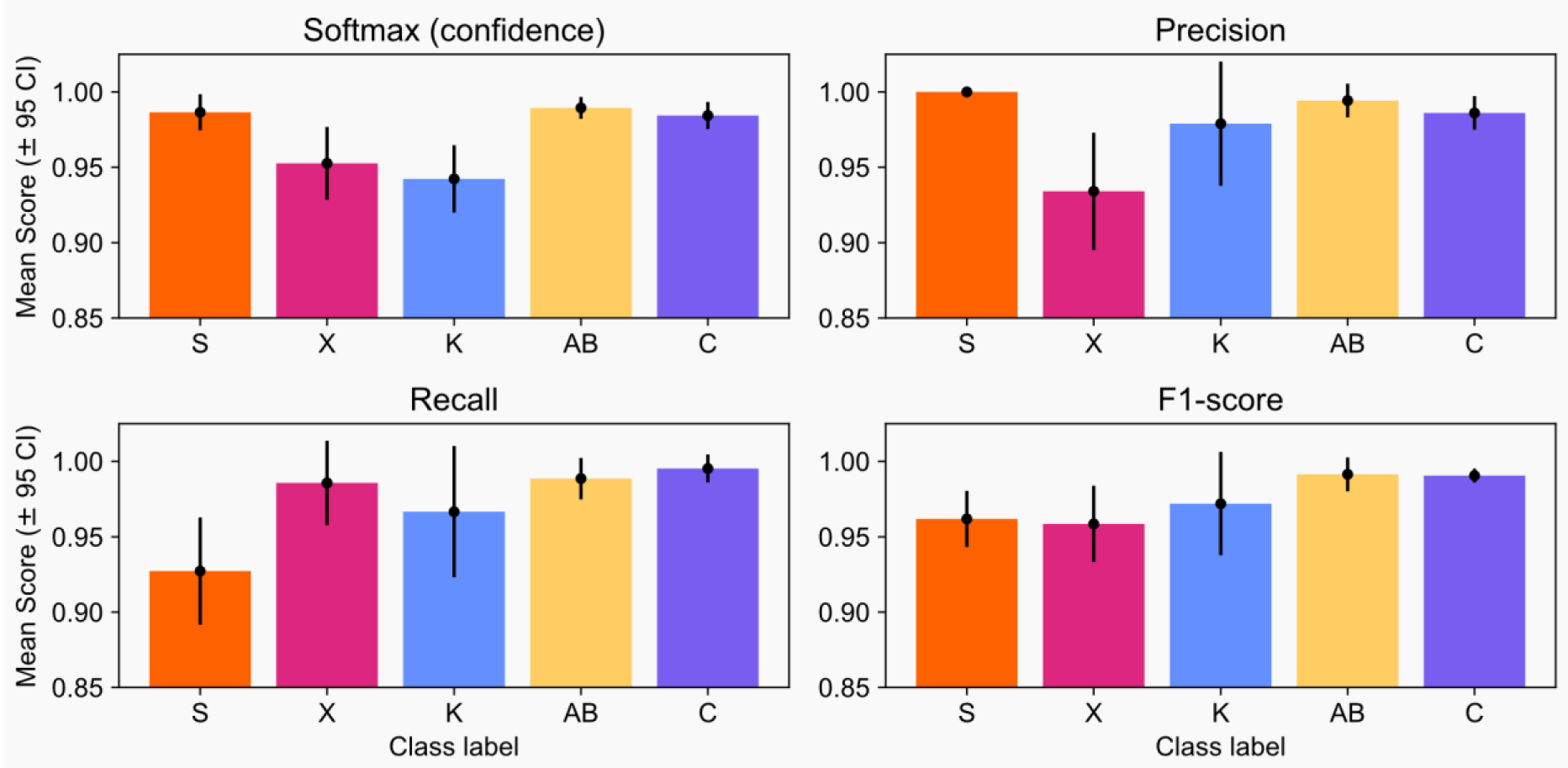
Softmax (confidence) and performance scores for class predictions. Classes are organised in order of increasing abundance (left to right).

#### 5.1.2. Visualize predictions

Another important component of model evaluation is to visualize the predicted class alongside the image, through the lens of a domain expert such as a benthic ecologist. Interrogating model decisions in this way can reveal patterns, error types, or limitations that are not apparent from the performance metrics alone. These insights may help explain misclassifications or, conversely, accurate class predictions. It also serves as an additional quality control check of the manual annotations.

A good approach is to group misclassified images by the annotator-assigned class and the predicted class and identify common, or unique, features that may cause confusion (Vega *et al*., 2024b). These features may correspond to technical characteristics, image quality or biological variation (see Section 2.1). More objective methods, such as Grad-Cam (Selvaraju *et al*., 2017), exist to identify which parts of the image the model weighs most heavily when assigning a class. Such tools can provide insight into error causes and guide improvements in the training data. However, these methods are outside the scope of this paper, largely due to their implementation complexity.

**Example 5.1.2 : Visual Analysis**

In our study, YOLO V.8m made limited misclassifications across the *k*-folds from which to learn, on average 2.2 mistakes per *k*-fold (total=11 across 7 unique images). Intuitively, the model was significantly less confident in the assigned class for these misclassified images, with an average confidence of 0.63 *(95% CI=* ±0.08*; SD=* 0.13*; n=*11), compared to 0.98 *(95% CI=* ±0.005*; SD=* 0.06*; n=*589) for those classified correctly. Examining the misclassified instances, many errors appear linked to features being common to multiple classes. In Figure 9(a) for example, an image of biotope K is wrongly classified as X, in which the dominant substrate appears similar, see *Figure 9(b)* for example. There are also some cases where intra-class variation may have led to misclassification for example Figure 9(c) & <u>9</u>(e) which have been incorrectly identified in multiple *k*-fold models. These appear less typical of their respective training images, Figures 9(d) & <u>9</u>(f), in terms of technical features such as lighting and perspective and ecological features such as substrate and turbidity.

#### 5.1.3. Assess practicality

Before deployment, it is important to decide whether the ML pipeline is acceptable in terms of usability, speed and effectiveness in the task it was set up for. Establishing a hard target for these is difficult as there is no community-approved, standardized or objective criteria that will suit every use case. Researchers must ultimately rely on their own judgment. Fundamentally, the use of ML is to provide some advantage over more laborious methods to justify the investment in skill hardware acquisition and development time. Several practical questions can also help guide this decision. Are experts satisfied with the performance metrics and how do these compare to the manual annotations if measured? How robust are the results under different conditions, for example different datasets (a non-trivial problem)? Is the task completed significantly faster? Given the resource requirements, for example hardware or data and human expertise, is the approach feasible and can it be scaled up? Establishing set criteria for the task will help to ascertain if the model is ready for real-world usage. Note that the answers to these questions could encourage a return to prior stages in the ML pipeline (Figure 1). However, this should be approached cautiously so as to avoid data leakage, in which the test data are used to guide adjustments to the ML pipeline, leading to biased improvements on test data. Instead you should aim to improve the generalizable performance without relying on test-set results, with cross-validation for example. If possible, it may even be prudent to use a novel test data set.

**Figure 9.**
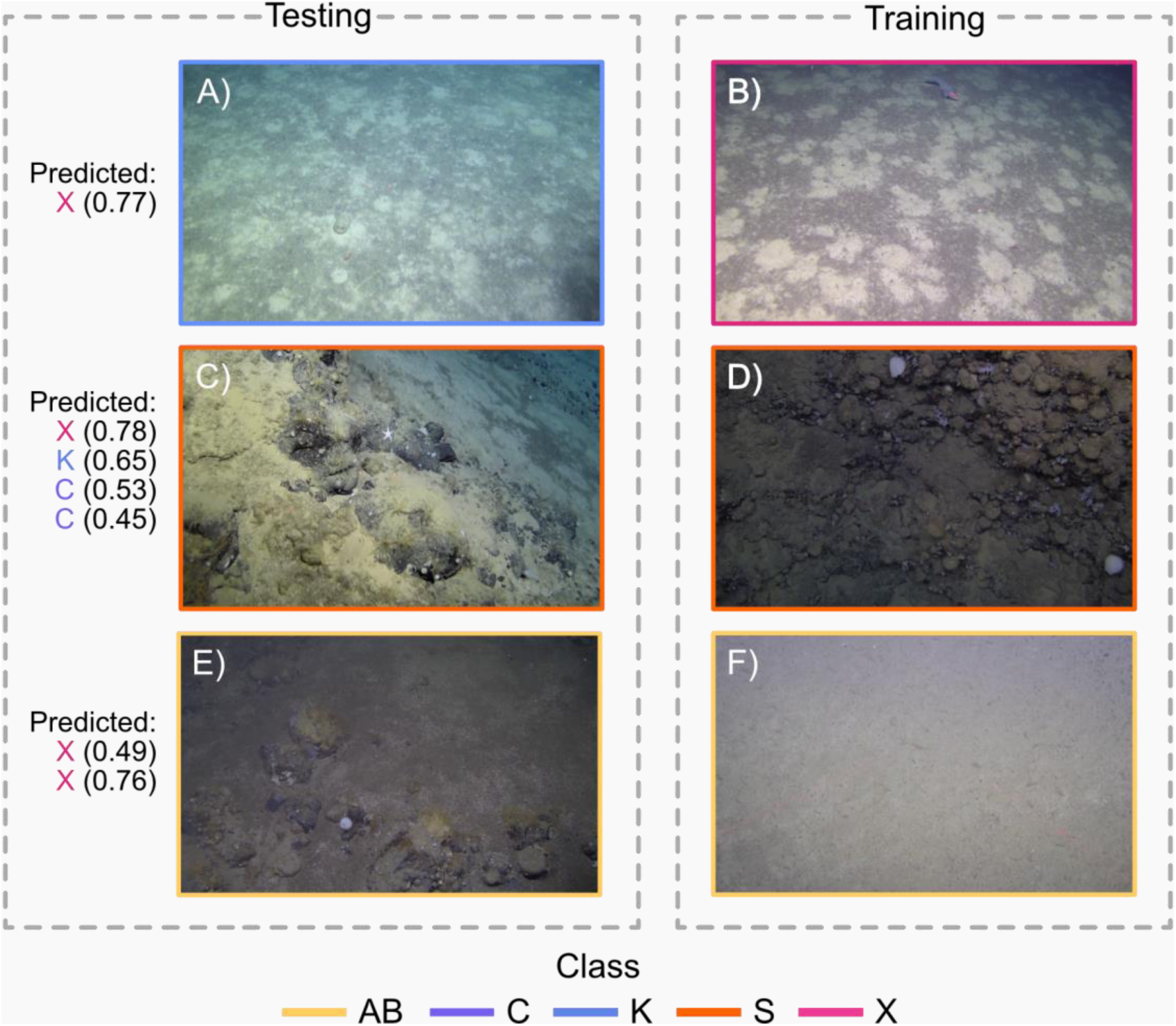
Examples of misclassified test images from all k-folds are shown in panels A, C & E, with predicted class labels and model confidence score in parentheses. The true class of each image is indicated by the border colour. Note that images in panels C & E were misclassified multiple times. For comparison, panels B, D & F display training images that either (1) share similar features but belong to a different class (A vs. B) or (2) differ in appearance but belong to the same class (C vs. D, E vs. F).

### 5.2 Save and deploy

In its simplest form model deployment means using the trained model to generate predictions on new images or video frames without subsequently updating the weights. Thus, saving the model avoids having to re-train every time. ML models can be saved in various formats which include the entire model (e.g. a .pkl file) or just the trained parameters (e.g. a .pt file). These can also be archived and shared on public model repositories such as Hugging Face (Hugging Face, 2025), Model Zoo (Model Zoo, 2025) or Roboflow (Roboflow, 2025). Both Hugging Face and Roboflow also offer options for private storage.

Model deployment can typically be performed on the same hardware used in training, provided the new data can be accessed by the system. The main limiting factors are data storage solutions (local hard-drive vs. cloud) and the type of imagery data (still images vs. video), rather than the modeling itself. This step can become a bottleneck since the imagery must be transferred to the machine running the model—via cable, network, or motherboard connections (the fastest). Transfer speeds may be significantly slower if either the data or the virtual machine resides in the cloud, especially for video data, which is substantially larger than still images. Note that, like in testing, the raw data must also be formatted consistent with the training procedure before any inference takes place (see Section 4.3).

As inference requires far less memory than training, it may even be feasible without a GPU within a practical time frame, depending on the architecture. Reduced computational demands mean models can be deployed on a wide range of hardware platforms and make real-time inference more accessible.

In general, there are some basic principles that can save time and effort when implementing automated image classification at large scales and over long periods of time. As with the efficient management of model-produced data, success requires careful planning and organisation. This is embodied in *MLOps*, or Machine Learning Operations, a set of practices, tools and processes used to deploy, monitor and maintain ML models (Kreuzberger, Kühl and Hirschl, 2023). Effective MLOps often determine whether an ML project successfully transitions from research into a sustained real-world application (Cooper, 2024). Broadly speaking, building a pipeline that is easily re-usable for another project or by other users can be just as (if not more) challenging than building the workflow in the first place. For long-running projects that will likely involve multiple users, considering ergonomy and long- term maintenance helps ensure the workflow gets used and actually saves time producing and delivering the data used to research the ecology. This can mean spending more time to keep the code tidy, commented, and modular so it can evolve easily.

Following any kind of inference, either during testing or wider deployment, model predictions should be exported. Typically this is saved as a table (e.g. a .csv file) with, at minimum, the image name and assigned class, although additional fields such as the softmax score can also be included. These outputs can then be used for further ecological analysis, including estimates of abundance, diversity and richness as well as community analysis. When combined with contextual environmental data from the ground-truth, for example geographic and bathymetric position, substrate type and water mass structure, this can also indicate distribution drivers and will support mapping, monitoring and marine spatial planning efforts.

The necessity of monitoring model performance throughout deployment should not be overlooked, as it is common to observe mismatches between training performances and field tests due to domain shift (e.g. new camera, different time of year with different water conditions and different communities from a different bioregion). Without regular checks to verify model predictions are in line with expectations of domain specialists, the models could feed erroneous information into the deliverables. Implementing a quality control procedure (as an extension of Section 5.1.2) and being ready to retrain the model when performances drop can alleviate these risks (Beery, Van Horn and Perona, 2018; H. Doig, O. Pizarro, and S. B. Williams, 2023).

**Example 5.2 (a) : Model Export**

A final biotope model (*k*-fold=3) was saved as a *.pt* file. Note that Ultralytics supports a range of other shareable formats for use within Pytorch and Tensorflow (Ultralytics, 2025b). The associated predictions from the testing phase were exported as a *.csv* and merged with the environmental ground truth (latitude, longitude and depth), generating the ecological knowledge for the final deliverable, in this case a distribution map (Section 6).

## 6. Deliverable: what ecological insights or outputs should the ML deliver?

The final deliverables from an ML application are project-specific but ultimately fall into two categories: tangible outputs and analytical insights. Like the objectives (Section 1), clearly defining both types at the project outset ensures that the methodology and analyses are aligned with the intended ecological outcomes or objectives.

Tangible outputs may consist of the trained model itself and the data products it generates, such as predictions on novel independent datasets, typically presented in a reprocessed and contextualized form alongside relevant metadata or complementary sensor data (e.g., occurrence records, diversity or abundance estimates, habitat change trajectories). Analytical insights comprise the ecological interpretation of what these outputs show (e.g. environmental drivers, spatial and temporal trends). They can also include quality assessment with uncertainty estimation for example, in which model confidence and potential errors are quantified; a particular advantage over manual analysis.

Reproducibility is a key aspect of both categories and should be maximised in all deliverables, as it underpins their comparability, standardization, and accessibility. In the context of this paper for example, the ML approach enables truly comparable seabed sampling, yielding standardized annotations and improving data accessibility to researchers and policy-makers (Jackett *et al*., 2023; Game, Thompson and Finlayson, 2024). Unlike the transfer of taxonomic expertise, sharing models is technically straightforward, and many frameworks exist for freely depositing and accessing them (Section 5.2). Such practices support FAIR (Findable, Accessible, Interoperable, Reuseable) principles (Wilkinson *et al*., 2016) and build trust in ecological analyses and biodiversity monitoring.

**Example 6 : Ecological Deliverable**

In this study, the end product is a distribution map (Figure 10) to visualize the predicted biotope classes with other spatially explicit data, enabling rapid identification of any ecological patterns. Since the aim here is to demonstrate the capability of the ML approach, rather than to provide a full ecological interpretation, refer to (Meyer *et al*., 2023) for a detailed ecological analysis of the ground-truth data.

The automated classification approach presented supports consistent analysis and mapping of these biotopes, and seabed communities more widely, facilitating the implementation of a standardized community classification system, such as “Natur i Norge” (NIN) in Norway (Halvorsen, Bryn and Erikstad, 2016) or EUNIS (Davies, Moss and Hill, 2004), as requested by policy makers (Howell *et al*., 2019b). By limiting biases, this approach also enables reliable detection of any distribution changes in benthic communities, which is crucial for monitoring dynamic ecosystems (Fraschetti *et al*., 2024).

**Figure 10.**
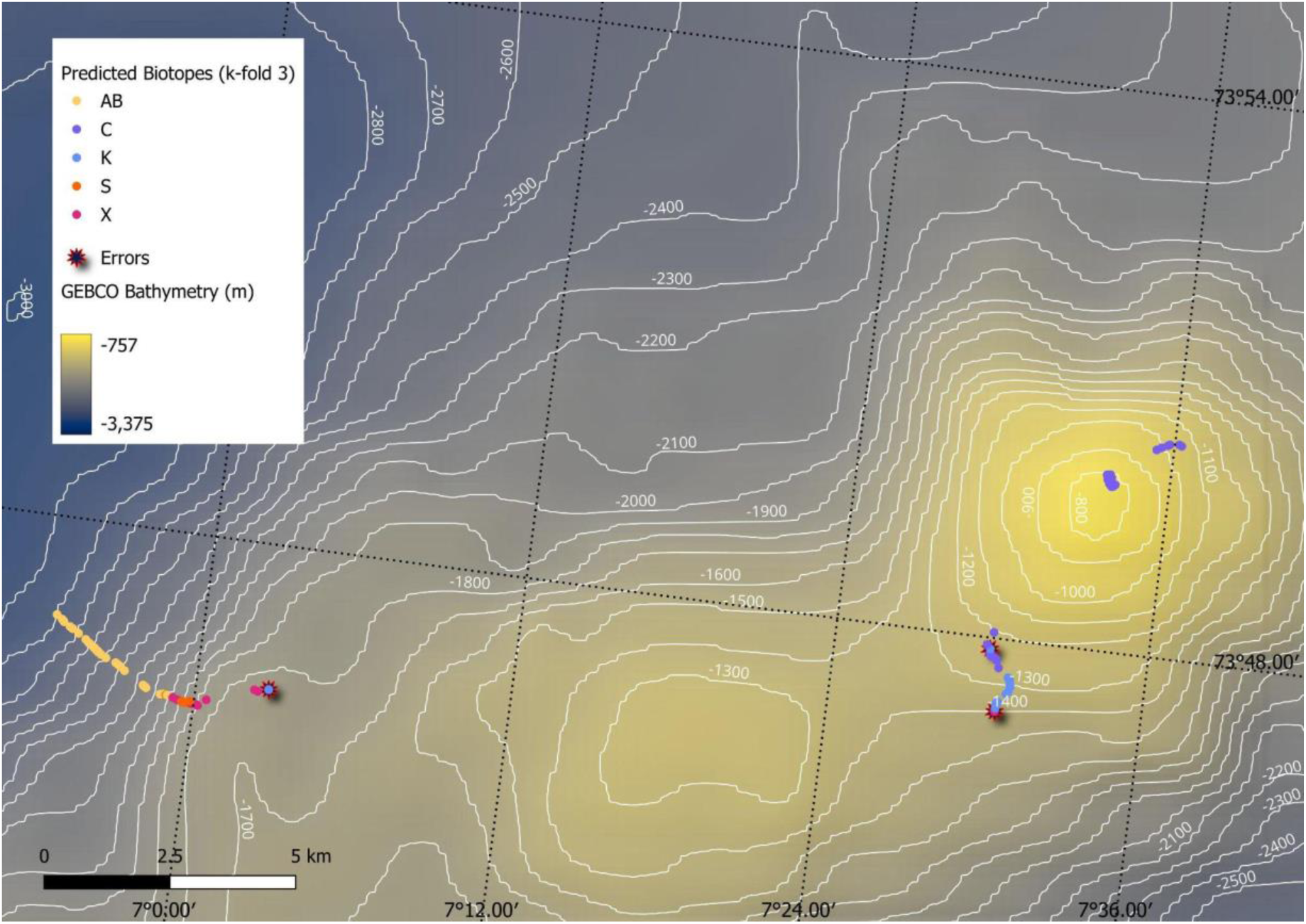
Map of the geographic distribution of the samples over the local bathymetry from GEBCO (GEBCO Bathymetric Compilation Group 2024, 2024) and 100m contours. Samples are colour coded according to their predicted classification in k-fold 3. The location of misclassified samples are also highlighted.*

## Conclusion

This study presented a simple, automated image classification workflow intended as a blueprint for approaching similar tasks, even if the specific model, computational framework, or ecological objective differs. The design is both modular and adaptable, with a flexible code base to expand upon. This flexibility is paramount for the wider adoption of such automated methods in ecology. It ensures continued relevance across applications and, since ML methods tend to evolve fast, allows users to integrate the latest and most efficient tools (e.g. ViTs) without redesigning the entire process. This not only saves time but preserves the expertise and familiarity acquired by the users and built within the rest of the workflow. Furthermore, it is unlikely that all stakeholders, in marine ecological research for example, will settle on the same set of tools in the future (e.g. annotation software, classification schemes and archiving infrastructures). Some degree of adaptation will, therefore, remain inevitable. Knowledge of the technical landscape is thus essential to select the best individual components for an application, integrate them effectively, and ensure compatibility in data formats and protocols. Sustained transdisciplinary collaboration and training will be key to this, ensuring both the automated tools and their associated data collection and processing remain aligned with the ecological objectives they are intended to serve.

Another overarching theme of this work was to improve accessibility for users with limited CV and ML experience, including, but not limited to, benthic ecologists. Using examples from the biotope case study, we provide users with enough context to facilitate adjustment to their own needs. More importantly, we highlight key *levers* for users to tune and optimize the model to their own application and demonstrate how these might influence the model results and usability.

For automated image analysis to deliver on its promises, users must not only explore its ability to emulate their routine outputs from manual analysis, but also commit to better data usage and management practices, as well as acquisition and retention of informatics skills (Schoening *et al*., 2022; Fraschetti *et al*., 2024). An advanced understanding of the underlying processes (e.g. ecological) that a model seeks to capture and predict is also capital for evaluating its efficacy before and during deployment. Domain knowledge should always guide the interpretation of predictions and the assessment of whether they can be used with confidence. As with other modelling frameworks, these automated approaches should be viewed as a powerful tool to extend, not replace, human insight.

## Acknowledgements

The authors would like to thank members of the SEAS Programme (University of Bergen) and the HIBRAIN working group (Institute of Marine Research) for their feedback on the code associated with this project. We also wish to thank Dr. Heidi Kristina Meyer for providing access to the Schulz Bank dataset and for their valuable guidance on the associated methodology. This study also acknowledges the use of GitHub Copilot, ChatGPT-4 and Le Chat for improving the efficiency of code development and documentation, as well as ChatGPT-4 and Perplexity AI for supporting intuition checks and grammar. C.A.G received funding from the European Union’s Horizon 2020 Framework Programme for Research and Innovation under the Marie Skłodowska-Curie grant agreement No. 101034309. N.P was supported by the MAREANO programme through the Institute of Marine Research, Norway. K.L.H was funded by NERC project UKRI041 “DEAL: DEcentrAlised Learning for automated image analysis and biodiversity monitoring”.

## Author contribution

C.A.G, N.P & K.L.H conceived the ideas for this work. C.A.G & N.P led the writing of the manuscript and literature review. C.A.G & N.P were responsible for data curation and designed the case study methodology and supporting code. C.A.G conducted the formal analysis and investigation of the case study. C.A.G, N.P & K.L.H designed the workflow and associated paper structure. Visualizations were produced by C.A.G. Mapping was conducted by N.P. All authors contributed critically to the drafts and gave final approval for publication.

## Data availability statement

The data used in this study were previously published by Meyer *et al*., (2022) and are publicly available at https://doi.pangaea.de/10.1594/PANGAEA.949920. To improve computational efficiency, a copy of the image data and biotope labels has also been made available at: https://huggingface.co/datasets/CGame1/schulz_bank_biotopes. All associated code (and chosen model) is publicly accessible at: https://cgame1.github.io/Img_classificaton_guide/).

